# Complete Enzyme Clustering Enhances Coenzyme Q Biosynthesis via Substrate Channeling

**DOI:** 10.1101/2025.05.24.655883

**Authors:** Dianzhuo Wang, Andrea Gottinger, Jio Jeong, Callum R. Nicoll, Junlang Liu, Tereza Kadavá, Domiziana Cecchini, Marco Malatesta, Albert J.R. Heck, Andrea Mattevi, Eugene I. Shakhnovich

**Author notes:** These authors contributed equally (Dianzhuo Wang, Andrea Gottinger, Jio Jeong, Callum R. Nicoll).

## Abstract

Metabolons - transient assemblies of sequential metabolic enzymes - facilitate the reactions of multi-step metabolic pathways, yet, how they mechanistically bolster metabolic flux remains unknown. Here, we investigate the molecular determinants of metabolon formation in coenzyme Q (CoQ) biosynthesis using coarse-grained molecular dynamics simulations and biochemical experiments. We show that the COQ metabolon forms at the critical region of a phase transition, where both metabolon clustering and metabolic flux exhibit coordinated sigmoidal responses to changes in protein-protein interaction strength. These complete metabolons enable substrate channeling between sequential enzymes, leading to a crucial enhancement of CoQ production efficiency. Selectively disrupting protein-protein interactions and randomly shuffling the interaction network demonstrate that protein-proximity rather than fine structure of the metabolon clusters is imperative for substrate channeling. Grounded in both experiment and simulation, these findings provide a framework for understanding the organization and function of metabolons across diverse metabolic pathways.

## Introduction

Protein-protein interactions affect the kinetics and dynamics of biological processes, fundamentally shaping cellular function.^1,2,3,4,5^ An example of this is seen in metabolic pathways, where cells assemble enzyme complexes known as metabolons - dynamic, multi-enzyme structures that enhance pathway efficiency and maintain metabolic homeostasis.^6,7,8^ Metabolons have been identified in both primary and secondary metabolism, organizing pathways such as the TCA cycle, *de novo* purine, flavonoid, and dhurrin biosynthesis.^9,10,11,12,13^ These supramolecular assemblies have been proposed to facilitate substrate channeling, which involves the swift transfer of intermediates between consecutive enzymes without the loss of metabolic productivity through diffusion into bulk solution.^8,14^ Metabolons thereby enhance metabolic flux, a principle that has been actively harnessed in synthetic biology. In metabolic engineering, scaffolds designed to spatially organize enzymes have successfully accelerated the biosynthesis of valuable metabolites, demonstrating the potential of rationally designed protein-protein interactions to optimize metabolic processes.^15,16,17,18,19^

Various theoretical works have provided insights into how enzyme clustering influences metabolic flux.^20,21,22,23^ Castellana *et al*. employed a reaction-diffusion model to study a two-step pathway metabolon in purine biosynthesis.^24^ Ranganathan *et al*. applied Langevin dynamic simulations of a patchy-particle enzyme model, demonstrating that enzyme clustering is particularly beneficial for reaction-limited enzymes.^25^ Other works have incorporated modeling or simulations to understand the physical origin of flux enhancement in artificial metabolons, synthetically designed multi-enzyme clusters that mimic natural metabolic channeling.^26,27,28,29,30,31^ Despite these advances and the growing recognition of natural and artificial metabolons, many questions remain regarding the molecular determinants of metabolon structure, the functional importance of specific protein-protein interactions, and how their architecture contributes to metabolic efficiency. The coenzyme Q biosynthetic pathway provides an ideal system to investigate such questions.

Coenzyme Q, also known as CoQ or ubiquinone, is the most hydrophobic cofactor in the cell, required for several essential functions. Widely known as an electron shuttle in the oxidative phosphorylation electron transport chain, the role of CoQ more recently been described in extra mitochondrial cellular processes, such as the anti-ferroptotic pathway, heavily implicated in cancer cell survival.^32,33^ CoQ is composed of a redox active polar head group and a poly-isoprenoid tail, whose length varies among species. In eukaryotes, the last steps of its biosynthesis take place in the mitochondrial matrix by means of several COQ proteins (Fig. 1a). COQ1 (PDSS1-PDSS2 tetramer) catalyzes poly-isoprene tail formation from activated isopentenyl-diphosphate units derived from the cytosolic mevalonate pathway, whilst COQ2 links the activated poly-prenyl-diphosphate tail to the C_3_ position of *para*-hydroxybenzoic acid obtained from cytosolic tyrosine catabolism.^34,35,36,37^ The benzoate polar head is then extensively modified throughout hydroxylation, O- or C-methylation and decarboxylation reactions. Such an elaborate series of decorations is performed by COQ3, 4, 5, 6, 7 and 9, forming an evolutionary conserved protein assembly referred to as the COQ metabolon, COQ synthome or complex Q.^38,39,40^ By employing stable ancestral tetrapod COQ proteins and soluble mono-prenylated biosynthetic intermediates, we recently characterized several unclear steps in the CoQ biosynthetic pathway (Fig. 1b).^41,42^ Moreover, COQ8B activity was shown to promote metabolon formation and channeling of intermediates.^41^ However, the mechanism behind channeling and the importance of cluster formation for metabolic flux remains unclear.

**Figure 1.**
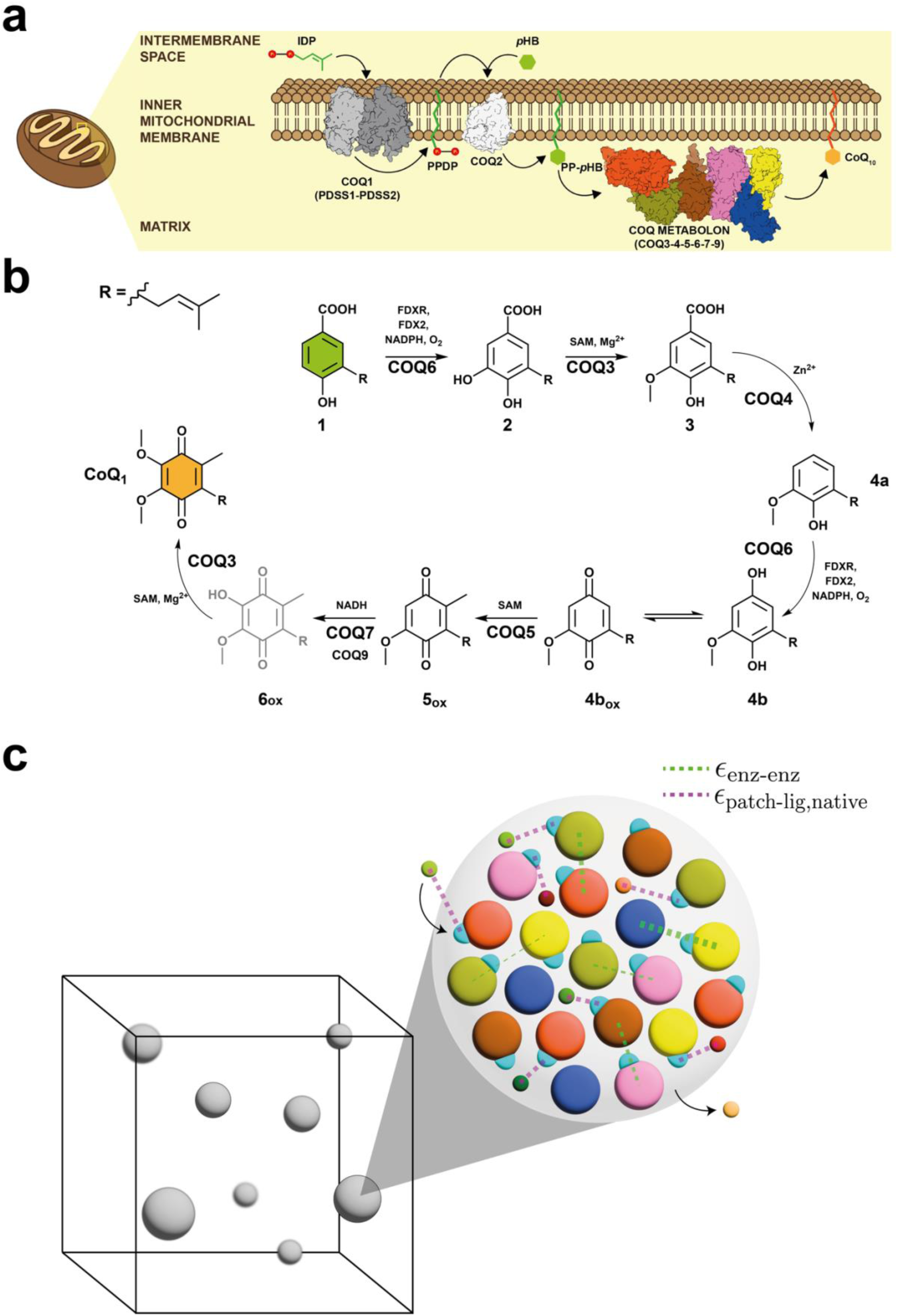
Coenzyme Q biosynthesis occurs in the mitochondrial matrix by means of the COQ metabolon. **a.** Poly-prenyl tail polymerization and linkage to the precursor head group occurs in the inner mitochondrial membrane whereas downstream head-group decorations are carried out by the COQ metabolon on the membrane’s matrix side. The precursor molecule is represented in green, CoQ in orange, enzymes that are not part of the COQ metabolon in gray scale, and COQ metabolon proteins are colored. pHB is p-hydroxybenzoate, IDP is isopentenyl-diphosphate, PPDP is isopentenyl-diphosphate. **b.** Reaction scheme for the reconstructed mammalian CoQ biosynthetic pathway. Especially relevant for this work, COQ6 is a class A monooxygenase that is responsible for both C_5_ and C_1_ hydroxylations and requires a protein-based electron donor - ferredoxin 2 (FDX2) and ferredoxin reductase (FDXR) - for its activation.^41,42^ COQ4 is Zn^2+^-dependent decarboxylase, responsible for C_1_ decarboxylation.^40–42^ **6_ox_** is the only biosynthetic intermediate not available for experimental work, and shown in transparency. **c.** Graphical representation of metabolon simulation showing the coarse-grained model used for computational studies. Gray colored spheres represent enzyme clusters that are scattered across the simulation box. In each metabolon, large colored spheres represent different COQ proteins (COQ3-7,9) with small blue patches indicating active sites. The interactions between enzymes (ε_enz-enz_) are shown by green dotted lines, while interactions between active sites and their native substrates (ε_patch-lig,native_) are indicated by pink dotted lines.

Here we employ both experimental biophysical analyses and coarse-grained molecular dynamics simulations incorporating experimentally measured protein-protein interaction strengths (Fig. 1c), to demonstrate that clustering of the full complement of COQ proteins (including COQ3-7 and COQ9) enables substrate channeling and dramatically enhances metabolic flux in the CoQ biosynthetic pathway. Our findings reveal that the COQ metabolon operates at the critical region of a phase-like transition, where small changes in protein-protein interaction strength can affect both enzyme clustering and metabolic output. We further show that the experimentally observed COQ interaction network is selected to promote the formation of complete enzyme clusters. We validate these computational predictions through targeted PEGylation experiments that selectively disrupt specific protein-protein interactions, demonstrating that metabolon completeness, rather than fine structure, is essential for efficient substrate channeling. These findings provide mechanistic insights into how cells achieve efficient multi-step biosynthesis through the spatial organization of metabolic enzymes.

## Results

### The COQ metabolon is held by an intricate and predominantly loose set of interactions

To precisely gauge the size of the metabolon and aid the molecular simulations, we used Size-Exclusion Chromatography coupled to Multi-Angle Light Scattering (SEC-MALS). We pooled together COQ3-7 and COQ9 (a total of six proteins), responsible for enzymatic activities required for head-group decorations, and observed co-elution of COQs in a soluble large molecular weight (MW) ensemble. Specifically, the elution profile showed a shoulder followed by two large peaks at MW of 461.4 (±1.2%), 265.6 (±2.2%) and 154.0 (±2.3%) kDa, respectively (Fig. 2a). With the cumulative weight of all COQ proteins amounting to 186 kDa, these data suggests that several larger complexes are present with increased stoichiometry beyond 1:1. In agreement, when considering an assembly corresponding to each reaction step, in other words, COQ6 and COQ3 twice, and the remaining COQs once, the theoretical MW is 265 kDa (Fig. 1b). Indeed, bottom-up proteomics on the high molecular-weight species isolated with analytical size exclusion chromatography revealed that COQ6 and COQ3 have relatively high (∼double) abundance (Fig. 2b). Further analysis by dynamic light scattering (DLS) revealed the presence of large hydrodynamic radius entities (Fig. 2c, Supplementary Fig. 1). Although minimal, this species echoes the larger entities (>1 MDa) reported from *in vivo* yeast studies.^43^ Such higher order assemblies are likely to be mediated by weak interactions.

**Figure 2.**
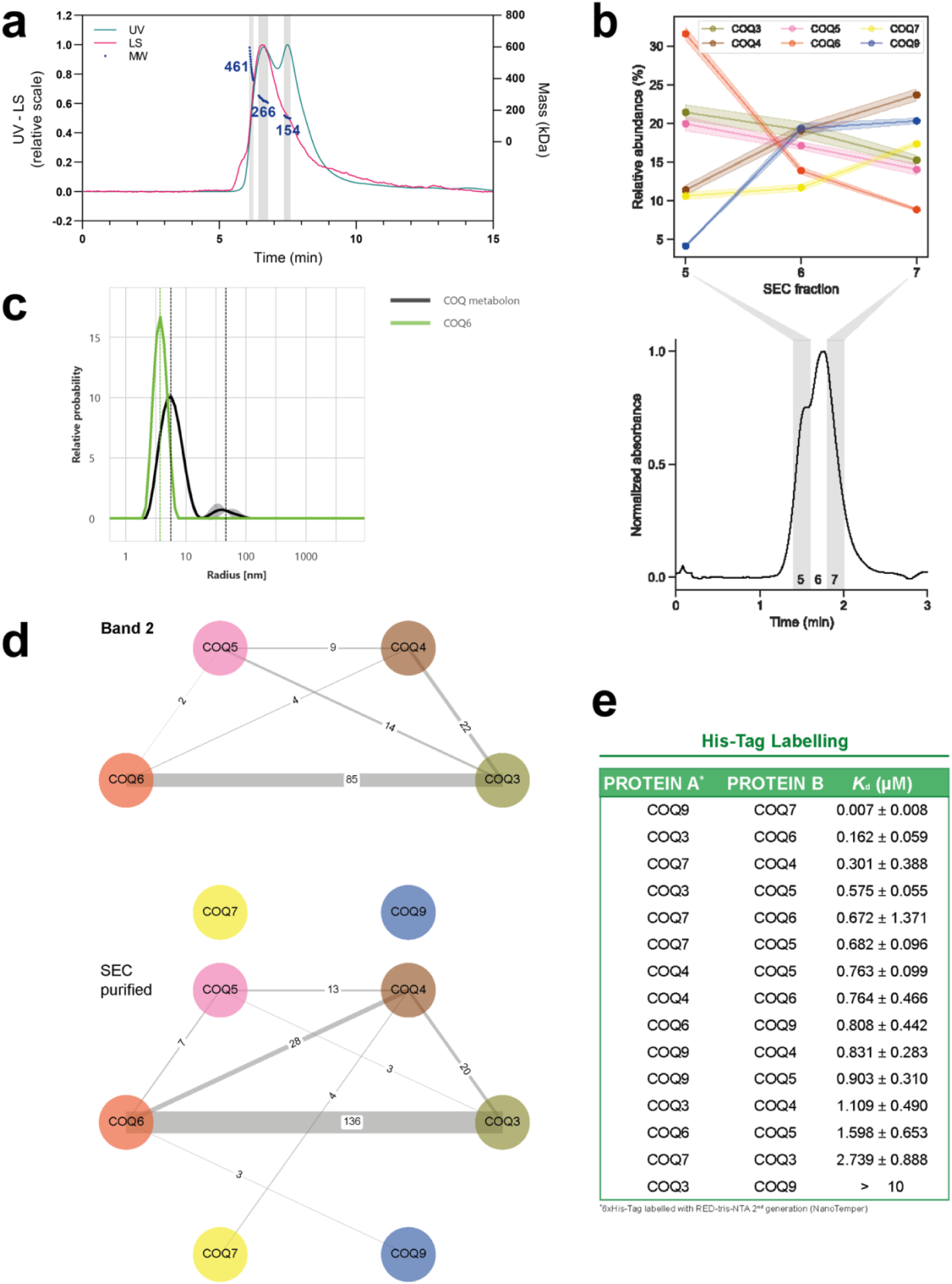
COQ proteins form high molecular weight species in solution, held together by an intricate interaction network. **a.** Size-exclusion chromatography coupled to multi-angle light scattering of the reconstructed COQ metabolon. UV at 280 nm signal is shown in green, light scattering (LS) signal in pink, and MW in blue. The analyzed peaks are highlighted by gray bands. **b.** Composition of COQ metabolon identified by bottom-up proteomics. Figure shows analytical size-exclusion chromatogram of COQ3-7 and 9, with highlighted fractions 5-7 (bottom). The upper panel shows the mean of normalized label-free intensities of COQ proteins across the fractions, as detected by bottom-up proteomics. The data are presented as mean of n=3 independent experiments, whose SD values are represented by the shaded areas. **c.** Dynamic light scattering of the COQ metabolon reveals large species. The panel overlays the profile of purified COQ6 (green) overlayed with the metabolon (black). Profiles of the other isolated COQ proteins are shown in Supplementary Fig. 1. The x-axis is in logarithmic scale. Data are shown as mean (line) and SD (shaded area) of n=4 independent measurements, each consisting of n=10 averaged acquisitions. Data are fitted with a size distribution model resulting in a hydrodynamic radius value of 5.64±0.13 nm for the main peak and of 46.10±12.84 nm for the second peak. **d.** Network plots obtained from (top) in-gel cross-linking mass spectrometry of the ∼240 kDa gel band, as annotated in Supplementary Fig. 15, and (bottom) in-solution cross-linking mass spectrometry of the size-exclusion chromatography (SEC)-purified COQ metabolon. Edges of the network are labelled with the number of unique cross-links (CSMs) found in the analysis. Data for gel bands 1 and 3 are provided in Supplementary Fig. 2. **e.** Pairwise affinity measurements of protein-protein interactions among COQs ranked based on dissociation constant values from tightest to looser interactions. Data are reported as mean and SD of n=2 independent measurements. Data were obtained with microscale thermophoresis using His-tag labelling strategy and shown in Supplementary Fig. 4. Data generated using primary amines covalent labelling are shown in Supplementary Fig. 3c.

To establish the protein-protein interactions involved in the assembly of COQs we first employed cross-linking mass-spectrometry (XL-MS). Both in-solution and in-gel revealed that COQs assemble in the metabolon through an intricate network of pairwise interactions, mainly involving the COQ3-4-5-6 core (Fig. 2d, Supplementary Fig. 2).^44^ To further investigate the assembly of COQs into supramolecular species, we then explored the protein-protein interaction network underlying metabolon formation using micro-scale thermophoresis. Since this method requires the target protein to be conjugated with a fluorophore, we employed two orthogonal labeling approaches: covalent modification of primary amino groups and His-tag labeling through non-covalent interaction with Ni²⁺-nitrilotriacetic acid. Using lysine-labeled proteins, we were able to detect a subset of pairwise interactions (Supplementary Fig. 3), while His-tag labeling enabled the measurement of dissociation constants (K_d_) for all COQ pairs (Supplementary Fig. 4). These findings suggest that surface labeling may interfere with measurements and highlight the advantage of more conservative techniques like tag labeling. Nevertheless, for interactions measurable by both methods, the K_d_ values obtained were comparable, validating the reliability of the measurements (Fig. 2e, Supplementary Fig. 3c). The results revealed a complex protein-protein interaction network with average K_d_ values in the mid-nanomolar range. The strongest interactions were observed for the COQ7-COQ9 pair (7 nM), for which the independent structure was solved experimentally, and for the COQ3-COQ6 pair (162 nM), which is also supported by the XL-MS data (Fig. 2d, Supplementary Fig. 2).^45^ These pairs may serve as more stable cores for the assembly of other COQs via dynamic and weaker interactions, ultimately leading to the formation of higher-order assemblies.

### Enzyme Clustering Enables Substrate Channeling and Enhances Metabolic Flux

The experimentally measured COQ protein-protein affinities provided the basis for Langevin dynamics simulations whereby the interactions between enzymes (ε_enz-enz_; reflecting the range of experimentally measured K_d_ constants; see methods) are included in a coarse-grained model of the metabolon. The enzyme activities are modelled as the reaction probability of 0.00001 for the substrate being converted into the product (*k_cat,base_*: Fig. 1c).^46^ As the first critical observation, the simulations revealed that enzyme clustering dramatically enhances the metabolic flux through the CoQ biosynthetic pathway (Fig. 3a). When protein-protein interactions were abolished, we observed a substantially reduced production of all pathway intermediates and nearly complete elimination of the final CoQ product (Fig. 3a). This effect was most pronounced for later intermediates in the pathway, with compounds **4b**, **5_ox_**, and **6_ox_**showing particularly stark differences between the clustered and non-clustered systems.

**Figure 3.**
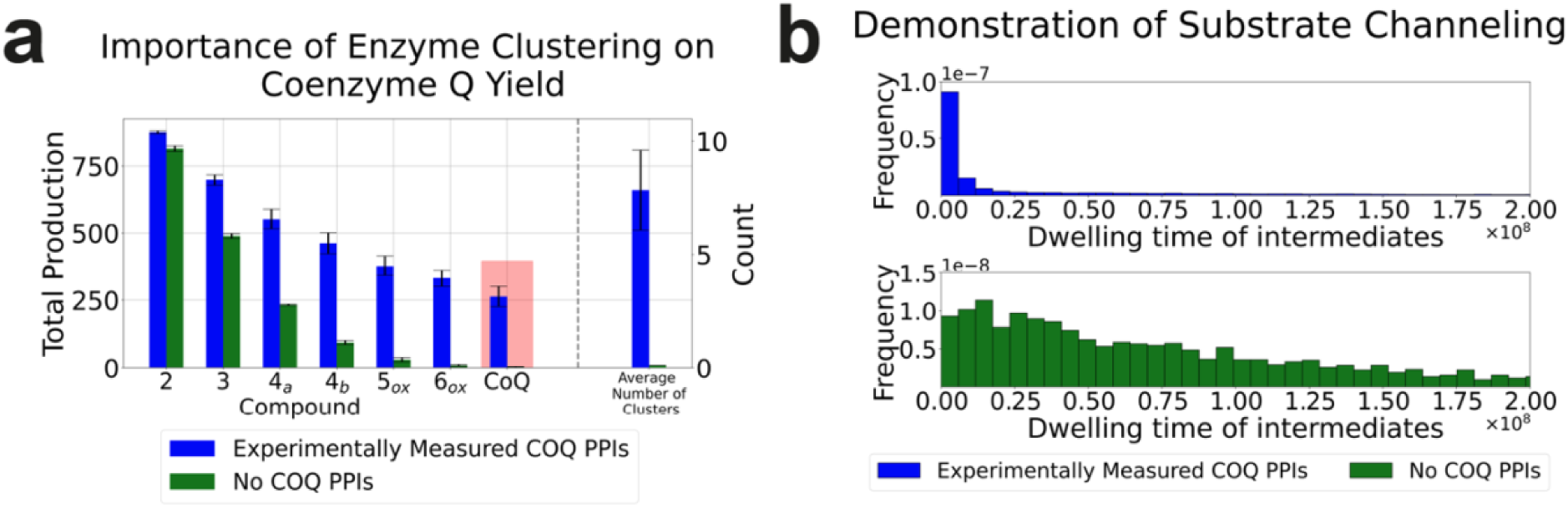
Enzyme clustering enhances metabolic flux through substrate channeling in the COQ metabolon. **a.** Comparison of intermediate and coenzyme Q (CoQ) yields between systems with experimentally measured COQ protein-protein interaction (PPI) strengths (blue bars) and systems without any protein-protein interactions (green bars). The data are presented as mean of n=6 and n=3 independent experiments respectively, whose SD values are represented by the error bars. **b.** Distribution of dwell times (the duration between consecutive reactions for each intermediate) in systems with experimentally measured COQ protein-protein interaction (blue) and no interactions (green). The left-shifted distribution in the clustered system (blue) indicates more rapid conversion of intermediates, demonstrating effective substrate channeling within the metabolon. Frequencies are normalized so that the values add up to 1, and dwell times are shown in units of simulation time steps.

To directly evaluate substrate channeling within enzyme clusters, we analyzed the temporal distribution of intermediate processing events. In systems with intact protein-protein interactions, the distribution of intermediate compounds’ dwell times (duration between consecutive reactions) was strongly skewed toward shorter intervals, with a sharp peak near zero and rapid decay. In contrast, systems lacking enzyme clustering showed a broader, more uniform distribution of dwell times (Fig. 3b). This left-shifted distribution in the clustered system indicates that intermediates are rapidly passed between enzymes within the same cluster rather than diffusing through bulk solution. These results quantitatively demonstrate that the spatial organization of COQ enzymes into clusters increases overall pathway flux by enabling efficient substrate channeling between consecutive enzymatic steps. This channeling mechanism helps explain how the COQ metabolon achieves efficient production of CoQ despite operating through multiple sequential reactions.

### The Experimental Protein-Protein Interaction Network Shows the Greatest Boost in Efficiency for Low Activity Enzymes

Having established the importance of enzyme clustering, we next investigated whether the specific pattern of the experimentally observed protein-protein interaction network is selected for metabolon function. Previous work by Ranganathan *et al*. suggested that metabolons might be particularly beneficial for enzymes with lower catalytic efficiencies, but the role of specific network topology remained unexplored.^25^ To test this, we developed a computational network randomization approach that preserved the distribution of interaction strengths while altering the network topology. This was achieved by rearranging the experimentally measured affinity values between different COQ protein pairs, effectively shuffling the network edges (Fig. 4a). The COQ7-COQ9 interaction was excluded from this randomization due to its known essential role in COQ7 catalytic activity.^41,45,47,48,49^ When comparing these randomized networks to the experimental network, we found that the latter consistently outperformed the former across all tested enzyme activities (Fig. 4b). Notably, the advantage of the experimental network was most pronounced at lower enzyme activities (Fig. 4c). When enzyme activity was set to 0.5 *k_cat,base_* (see *k_cat,base_* definition in Methods), the experimental network showed a two-fold advantage over randomized networks in CoQ production, whereas this advantage gradually decreased at higher activity of the enzymes, becoming almost undetectable at 10 *k_cat,base_*. This sensitivity of low-activity enzymes to enzyme clustering within the metabolon can be explained by considering where reactions predominantly occur. With low *k_cat_* values, only about 7% of reactions occur in bulk solution, with the vast majority taking place within enzyme clusters. As activity increases, the proportion of bulk reactions rises to 20% (Fig. 4c), reducing the system’s reliance on clustered reactions. Consequently, randomizing the interaction network has a more severe impact on low-activity enzymes, where disruption of the optimized cluster composition strongly reduces product formation. In contrast, higher-activity enzymes are more tolerant to network perturbations because their enhanced catalytic activity allows them to partially compensate through bulk-phase reactions, resulting in a smaller advantage for the experimental network topology.

**Figure 4.**
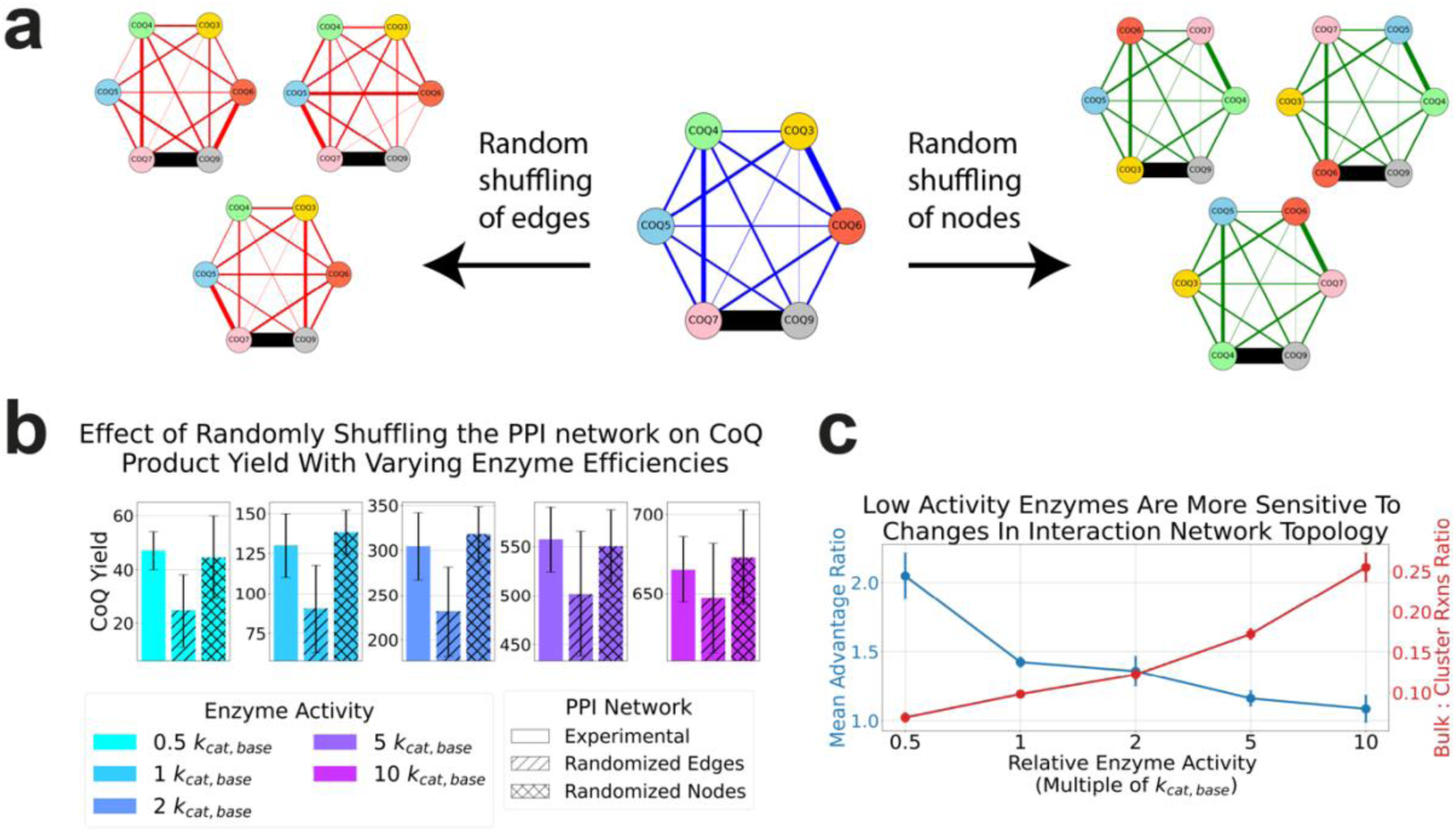
Impact of protein-protein interaction network topology on metabolon function and enzyme clustering. **a.** Schematic representation of the PPI network randomization process for both edges and nodes. The experimentally determined network (blue) shows interactions between COQ proteins with line thickness indicating interaction strength. The COQ7-COQ9 interaction (black line) was preserved during randomization of edges. The COQ9 node was fixed since it does not possess catalytic activity. Three representative examples of each randomized network are shown. **b.** Comparison of coenzyme Q yields between experimental (solid bars), randomized edges (hatched bars), and randomized nodes (cross hatched bars) PPI networks across different enzyme activities. The data are presented as mean of independent experiments (n=6 for experimental, n=12 for randomized edge networks, n=6 for randomized node networks), where error bars represent standard deviation. **c.** Analysis of network topology sensitivity across a range of enzyme activities. The advantage ratio (blue, left axis) shows the relative performance benefit of the experimental network over randomized edge networks in terms of final coenzyme Q yield. The data are presented as a mean of independent simulations (n=6 for experimental interaction network, n=12 for randomized edge network), whose SD values are represented by the error bars. See Yield Ratio Calculation in Methods. The bulk cluster ratio (red, right axis) indicates the ratio of reactions occurring in bulk solution to those within enzyme clusters. The data are presented as a mean of n=6 independent simulations of the experimental interaction network, whose SD values are represented by the error bars.

### Fine Structure is not Essential for COQ Metabolon Yield

To understand the mechanistic basis of the experimental network’s superior performance, we considered a hypothesis that the specific affinities might determine the fine structure of the metabolon by controlling the spatial arrangement of enzymes within clusters, akin to well-established complexes of proteins such as ribosomes or proteasomes. In this scenario, differences in spatial arrangement of proteins inside clusters would directly influence metabolic flux. To test this “fine structure” hypothesis, we performed a more stringent test by randomizing the nodes of the experimental protein-protein interaction network while preserving its topology (Fig. 4a).

Focusing on three different node-shuffled interaction networks, we analyzed the radial distribution function (RDF) of consecutive enzyme pairs - enzymes that catalyze sequential reactions in the pathway (Supplementary Fig. 5). The RDF analysis revealed drastically different distributions between the networks, showing that the spatial arrangement is affected by node randomization (Supplementary Fig. 6). If the specific spatial arrangement of consecutive enzymes was crucial, this perturbation would dramatically affect CoQ yield. However, node randomization had minimal impact on product formation (Fig. 4b, Supplementary Fig. 6), providing strong evidence against the fine structure hypothesis.

### Complete Enzyme Clusters Drive Metabolon Efficiency

We next considered the more basic hypothesis that network topology of PPI primarily functions to promote the formation of enzyme clusters that are complete, i.e. contain all metabolon enzymes. To test this hypothesis, we analyzed the relationship between CoQ production and cluster composition. We found a strong correlation (Spearman’s ρ = 0.89, P = 5.76e-09) between the final product yield and the proportion of enzymes that are members of complete clusters containing at least one copy of each required COQ enzyme (Fig. 5a, Supplementary Fig. 7). Furthermore, the experimental network consistently maintained a higher proportion of enzymes in complete clusters compared to randomized networks. (Fig. 5a)

**Figure 5.**
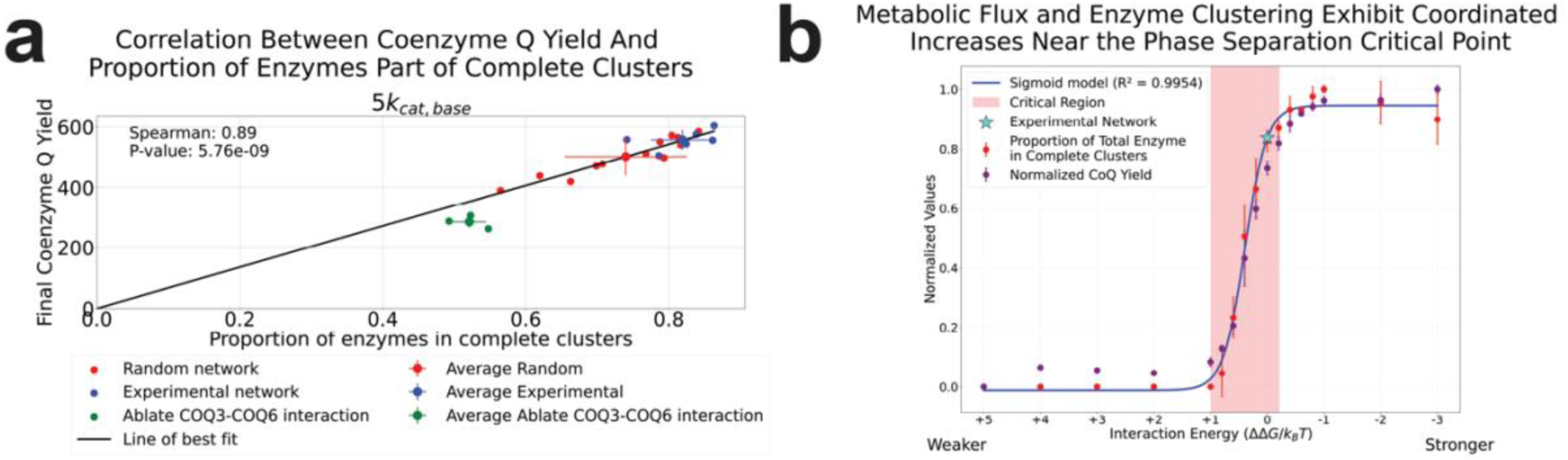
Complete Enzyme Clusters Drive Metabolon Efficiency. **a.** Correlation between coenzyme Q yield and the proportion of enzymes in complete clusters (containing at least one copy of COQ3-7). Each data point represents an individual simulation for experimental network (n=6, blue), randomized networks (n=12, red), and COQ3-COQ6 interaction ablated (n=3, green). The data points with the error bars represent the mean value across independent simulations of each condition, where the error bars represent the standard deviation. **b.** Metabolic flux and complete enzyme clustering exhibit coordinated increases near the phase separation critical point. Red points show proportion of total enzymes participating in complete clusters, while purple points show normalized CoQ yield. The data are presented as a mean of n=3 independent simulations for each interaction energy value, except for the experimental network at an interaction energy of 0, for which n=6. The error bars represent the standard deviation. Blue line represents the sigmoidal model fit (R² = 0.9954) to total enzymes participating in complete clusters, with the critical region highlighted in pink. Experimental network (⋆) is positioned within this critical region.

Having established the importance of complete clusters for metabolic output, we next investigated how the strength of protein-protein interactions influences cluster formation and metabolic efficiency. In Fig. 5b we varied the overall strength of PPI in the cluster by adding/subtracting a constant energy offset 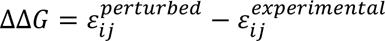 to each pair of interactions. This scan covers a broad range of interaction strengths from weak interactions where clusters are not formed, to strong interactions which lead to clustering. We observe that the efficiency of the enzymatic chain and the proportion of enzymes in complete clusters closely follows the sigmoidal curve (Fig. 5b). Notably, both transitions occurred at the same energy threshold, indicating a strong correlation between complete metabolons and metabolic output. Furthermore, we observe that the experimental network is located at the critical region of this phase transition, where the system exhibits highest sensitivity to changes in interaction. These results show that enzymatic efficiency of the whole chain is contingent on formation of the complete metabolon and that formation of the complete metabolon occurs as a phase transition akin to liquid-liquid phase separation.

To additionally validate the notion that completeness is the key factor, we systematically disrupted cluster formation by preventing individual COQ enzymes from participating in protein-protein interactions (Supplementary Fig. 8). This approach, which ensures the formation of exclusively incomplete clusters, completely abolished CoQ production for every COQ enzyme kept in isolation, suggesting that the experimentally observed network topology is above-all optimized to promote the formation of complete enzyme clusters necessary for efficient substrate channeling through the entire pathway (Fig. 6a).

**Figure 6.**
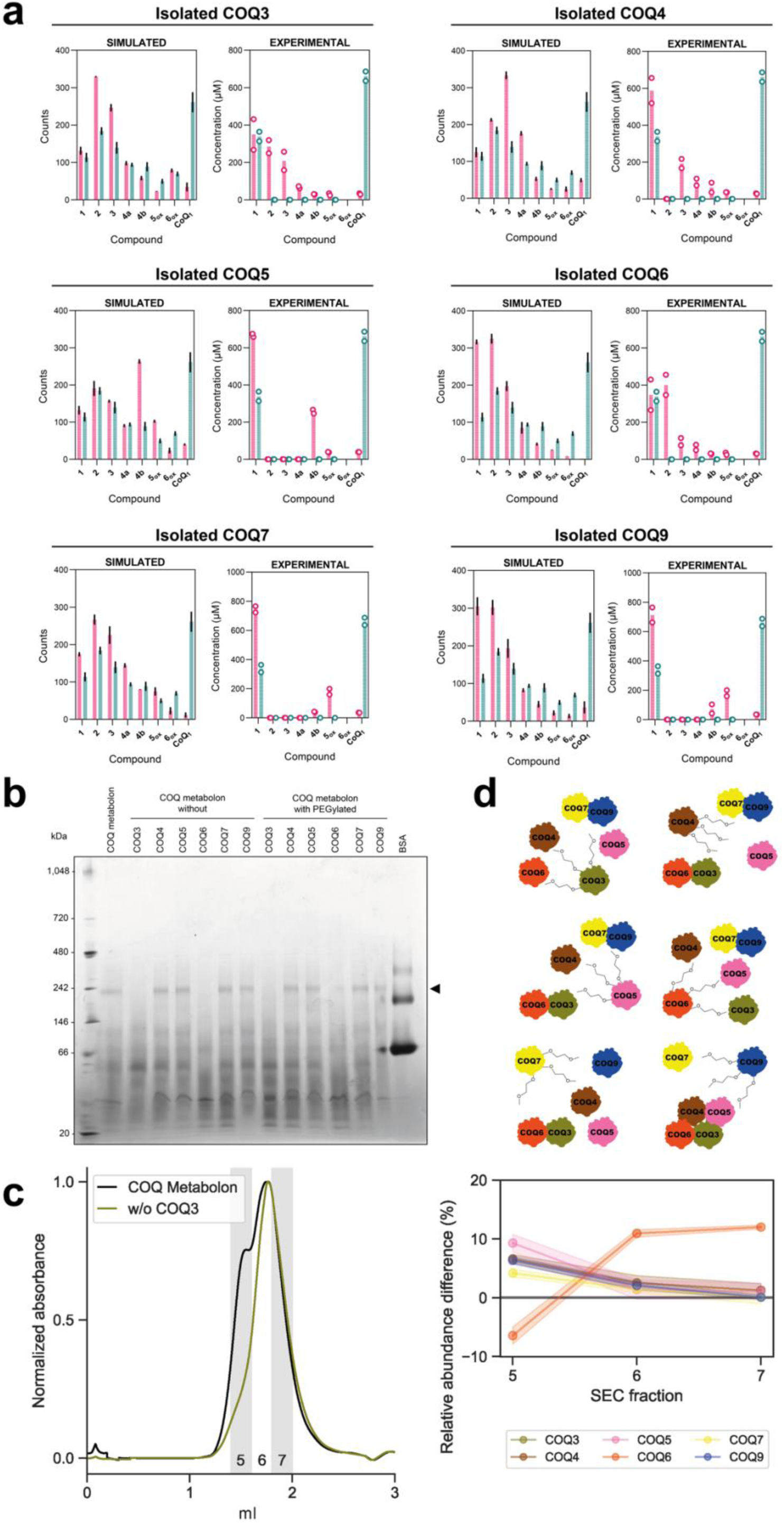
Disruption of the COQ metabolon interaction network drastically alters activity and topology. **a.** Pairwise comparisons of simulated and experimental COQ metabolon activity using isolated COQ proteins. Isolation (green) was carried out by ablating interactions for a given protein in simulation and by chemical PEGylation in experimental reactions. Controls with the intact metabolon are shown in green. Data shown in all panels are overlayed with the same control experiment. Simulation data are shown as mean and SD of n=3 independent replicates, while experimental data are shown as individual data points from n=2 independent measurements. **b.** Blue Native PAGE analysis of the intact COQ metabolon, with removed or PEGylated proteins (from left to right). The band at about 242 kDa, if present, was subjected to trypsinization and peptide mapping is marked by a black arrow ( Supplementary Table 1). The uncropped gel is reported in Supplementary Fig. 18. **c.** Comparison of intact COQ metabolon and deprived of COQ3. Figure shows size-exclusion chromatography chromatogram (left) of the COQ metabolon with (black) and without COQ3 (olive green). The right panel highlights compositional differences between incomplete and full COQ metabolon as detected by bottom-up proteomics. The data are presented as mean of n=3 independent experiments, whose SD values are represented by the shaded areas. Analyses of the metabolon deprived of the other COQ proteins are reported in Supplementary Fig. 13, corresponding SDS-PAGE gels in Supplementary Fig. 19. **d.** Topological models of the altered COQ after metabolon PEGylation fitting experimental data.

To test this computational prediction, we isolated individual COQ proteins from the rest of the assembly using chemical surface modification that disrupts protein-protein interactions one protein at a time. Given that lysine labelling impaired several protein-protein interactions in MST experiments (Supplementary Fig. 3c), we employed once more N-hydroxysuccinimide chemistry to modify COQ surfaces with a bulkier methoxy-PEG 5,000 moiety (Supplementary Fig. 9). Given that AlphaFold-3 predictions indicated an even distribution of lysine residues across the COQ surfaces (Supplementary Fig. 10), we hypothesized that the methoxy-PEG long chain could act as a steric spacer, obstructing direct protein-protein interactions. As a proof-of-concept we investigated the impact of PEGylation using a pull-down assay with His-COQ6 as bait (Supplementary Fig. 11). While unlabeled COQ3 co-eluted with COQ6, PEGylated COQ3 eluted in flow-through and wash, demonstrating that PEGylation disrupts the COQ3-COQ6 interaction. Next, we assessed enzymatic activity and confirmed that PEGylation preserves protein folding and enzymatic activity (Supplementary Fig. 12a). Notably, PEGylated COQ6 and COQ7 turned out to be two-fold more active than the unmodified proteins, aligning with reports of enhanced PEGylated enzyme stability and activity (Supplementary Fig.12b).^50^ With evidence that PEGylated COQs remain catalytically active as individual enzymes, we investigated their effect on the COQ metabolon in the presence of COQ8 that enhances its activity. Each PEGylated COQ was tested individually in an overnight reaction containing the other COQ proteins, cofactors, and precursor **1**. All biosynthetic intermediates, except for **6_ox_**, for which no analytical standard is available, and coenzyme Q_1_ were quantified using LC-MS (Fig. 1b, Fig. 6a). Each PEGylated protein could no longer effectively channel their substrate as we observed accumulation of its substrate and upstream intermediates, along with a roughly 15-fold reduction in CoQ yields (Fig. 6a). Previous work illustrated that COQ8, an atypical kinase, was imperative for efficient metabolon function and therefore included in the metabolon used for the experimental analysis.^41,51,52^ Here we did not scrutinize the precise molecular mechanisms by which COQ8 is able to boost metabolon activity, but the experiments with the PEGylated proteins suggest that COQ8 requires enzyme-enzyme interactions to be intact. These results corroborate the conclusions from simulations of the incomplete clusters and demonstrate that COQ metabolon completeness is necessary for substrate channeling through the entire pathway.

PEGylation of COQ3 and COQ6 had the strongest effect as it disrupted pathway channeling across the entire metabolon. Conversely, PEGylation of the other COQs did not impair channeling upstream of their catalyzed reaction (Fig. 6a). To explore the structural aspects of these results, we assayed the metabolon with native-PAGE after removing a protein or replacing it with a PEGylated COQ enzyme one at a time. Crucially, the band at an apparent mass of ∼240 kDa, corresponding to the metabolon, disappeared exclusively when COQ3 or COQ6 were either removed or PEGylated (Fig. 6b). Similarly, size-exclusion chromatography profiles and bottom-up proteomics analysis of the COQ metabolon deprived of either COQ3 or COQ6 showed the loss of the large species, whereas removal of other COQs did not result in significant changes (Fig. 6c, Supplementary Fig. 13). These results underscore the pivotal structural and functional role of COQ3 and COQ6 in the COQ metabolon. We conject that the COQ3-COQ6 bond acts as the cluster cornerstone, on which other COQs can assemble and posit that other metabolons possess similar assembling agents (Fig. 6d).

### Inactive COQ Proteins Preserve Metabolon Integrity and Channeling

To further examine the structure-function relationship between the COQ metabolon architecture and the optimization of metabolic flux, we reconstructed the metabolon using catalytically inactive COQs. The goal was to maintain a structurally intact metabolon that is enzymatically inactive for specific reactions. To achieve this, we utilized COQ6 without the addition of FDXR and FDX2, which are required for its activity (Fig. 1b), and employed the COQ4 H142A-H146A mutant, previously shown to be inactive due to disruption of the Zn^2+^ binding motif.^41^ For the experiments involving inactive COQ4, we initiated the reactions with both precursor **1** and intermediate **4a** to bypass its step in the pathway. In experiments with inactive COQ6, we bypassed its reactions by starting from intermediate compounds **2** and **4b** (Fig. 1b). The results showed higher yields compared to the PEGylation experiments and, more notably, evidence of intermediate channeling similar to the control experiments (Fig. 7a). These findings, combined with evidence that the metabolon is assembling regardless of the active site COQ4 mutation (Fig. 7a-c), suggest that the benefits conferred by the condensation of COQ proteins into a metabolon are predominantly associated with its structural integrity.

**Figure 7.**
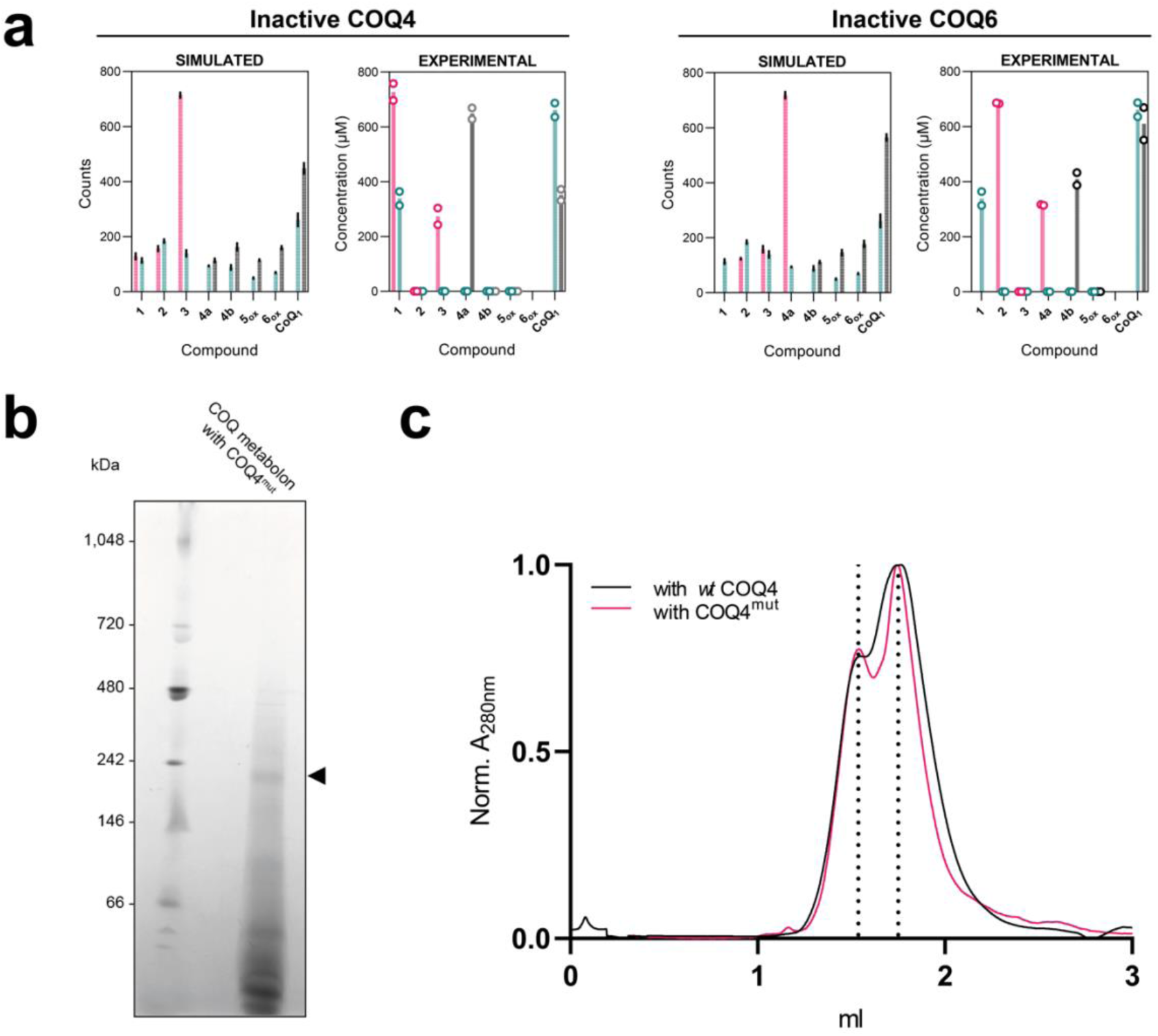
Inactive COQs preserve the COQ metabolon architecture. **a.** Pairwise comparisons of simulated and experimental COQ metabolon activity using inactivated COQ4 and COQ6. COQ4 H142A-H146A mutant was used to inactivate COQ4, whereas COQ6 was inactivated by removing the ferredoxin pair from the reaction. Control experiments with the intact metabolon are shown in green. The first half pathway prior to the inactivated step in pink, the second half pathway in black. Simulation data are shown as mean and SD of n=3 independent replicates, while experimental data are shown as individual data points from n=2 independent measurements. Compounds **2** and **4b** were added to the experiments with inactive COQ6 whilst **1** and **4a** were added to the metabolon with inactive COQ4 (Fig. 1b). **b.** Blue native PAGE analysis of the COQ metabolon including mutant H142A-H146A COQ4. The band at about 242 kDa subjected to trypsinization and peptide mapping is marked by a black arrow (Supplementary Table 2). The uncropped gel is reported in Supplementary Fig. 18. **c.** Analytical size-exclusion chromatography profiles of the COQ metabolon including H142A-H146A COQ4 mutant compared with the profile of the non-mutated (wt COQ4) metabolon, shown in pink and black respectively. The elution volume of the peaks in the control chromatogram are marked by dashed vertical lines.

## Discussion

Srere originally proposed that metabolons are supra-molecular complexes composed of sequential metabolic enzymes and cellular structural elements.^6^ While previous studies have examined the effect of enzyme proximity on metabolic flux in two-step pathways using reaction-diffusion models and numerical simulations^24^, few have explored longer pathways due to the increasing complexity with each additional step state.^20,23,25,26,28,53,54^ To broaden our understanding of intracellular organization of these key pathways and more specifically, the molecular determinants associated to metabolon formation and spatial enzyme organization, we employed a two-pronged approach merging experimental and computational methods to scrutinize the iconic system - the COQ metabolon - involved in mitochondrial CoQ biosynthesis. Our integrative approach, combining biophysical and computational analyses revealed several fundamental insights about metabolon function.

i. *Almost each COQ protein can interact with any other*, displaying K_d_ values between 100 nM and 10 μM, except for COQ7-COQ9 which interacted at around 7 nM and COQ3-COQ9 which did not show conclusive interaction (>10 μM). These somewhat mid-range protein-protein interactions are in line with previous literature that suggests these supramolecular complexes are loosely held together.^6,25,55,56^ Indeed, weakly interacting protein assemblies are easier to dismantle and reassemble as need arises, which has been found to be leveraged by the cell as a regulatory characteristic of both the purinosome and glycolytic metabolon.^57,58^ Their ability to assemble and accelerate final product synthesis in multi-step pathways in response to biological cues such as oxidative stress and cellular energy demands could provide a significant survival advantage.^59,60,61,62,63^
ii. *The benefits of metabolon formation become more pronounced along the reaction sequence*. The enzyme clustering into agglomerates facilitates the rapid processing of intermediates, providing similar benefits to direct channeling even without physical tunnels between enzyme active sites.^24^ Formation of early intermediates, such as **2**, showed minimal dependence on clustering, while later steps became increasingly reliant on proper enzyme organization with a complete loss of CoQ yield in non-clustered systems.^25^
iii. *Complete metabolon clustering, but not its fine structure, is pivotal for metabolic flux.* Despite the COQs being enzymatically competent as standalone systems, we observed that when clustering is disrupted, CoQ yields dropped significantly, in agreement with previous *in vivo* yeast studies.^48^ In both experiments and simulations, accumulation of the isolated enzyme’s substrate was observed. This observation suggests that substrate channeling up to the isolated enzyme is uninterrupted. One notable difference though is that in experiments, intermediates were undetected when there was efficient substrate channeling, whereas in simulations, even in the control with complete metabolons, intermediate compounds were always detectable in the system. This discrepancy underscores the inherent limitations of both approaches: the detection limit of compounds in small-scale reactions and the challenge of simulating timescales that accurately reflect experimental conditions. Despite these issues, both simulations and experiments demonstrate the importance of completeness in clusters. We propose that while the metabolon does not possess a rigid structure per se, it does however possess a minimal global structural hierarchy that is crucial for efficient molecular channeling facilitated by COQ8.^51,52^ Indeed, our study demonstrates that the COQ3-COQ6 interaction is imperative for the proper structural and functional organization for the COQ metabolon. The observation that complete metabolon formation is essential for efficient CoQ synthesis suggests that mutations affecting protein-protein interactions, not just catalytic activity, could lead to pathological CoQ deficiency.^64^ Nevertheless, it is important to acknowledge that our coarse-grained computational approach, while providing insights into the functional importance of enzyme clustering, necessarily simplifies certain aspects of metabolon structure and dynamics. The model treats enzymes as spherical particles with adhesive interaction sites, which may not capture more subtle structural features that could exist in the native cellular environment, such as specific binding orientations, conformational changes upon binding, or allosteric effects. These simplifications might underestimate the importance of specific protein-protein interfaces that could fine-tune substrate channeling efficiency. However, the consistency between our computational predictions and experimental results regarding the importance of cluster completeness over fine structure suggests that these potential structural nuances, while possibly present, are not essential for the core functional properties of the COQ metabolon. This finding highlights the robustness of the metabolon’s design, wherein proximity of multiple copies of the complete enzyme set appears to be the predominant determinant of functional efficiency rather than precise structural arrangements.
iv. *The specific pattern of protein-protein interactions appears to be evolutionarily selected.* The assembly of the COQ metabolon exhibits characteristics of a phase transition, with both enzyme clustering and metabolic output following coordinated sigmoidal transitions at the same interaction energy threshold. Remarkably, the experimental network appears to be located at this critical region of the phase transition. The positioning of the system at criticality may represent an evolutionary solution that balances the competing demands of CoQ output, and regulatory responsiveness.^65^ Randomly shuffling PPI consistently produced lower CoQ yields and lower degree of cluster completeness than the experimental network, with this effect becoming more pronounced for enzymes with lower catalytic activities. This suggests that this specific parameter regime of the PPI network has been selected for complete metabolon assembly, a prerequisite for substrate channeling and minimal dissipation of intermediates. Future work could investigate whether the critical phase transition behavior observed in the COQ metabolon is a common feature across other metabolic pathways, and whether cells actively regulate interaction strengths to modulate metabolic flux in response to changing conditions.

Ultimately, our findings have broad implications for the *de novo* design of synthetic metabolons in metabolic engineering and for the rational design of protein scaffolds and multi-enzyme fusion.^15,16,17,18,19^ Engineering protein-protein interactions to build artificial metabolons remains challenging, especially as pathway complexity increases. Inspired by Wang *et al*., who showed enhanced activity for an engineered two-bodied system with facing active sites,^19^ we hypothesize that artificial metabolons may exploit similar spatial arrangements to facilitate efficient catalysis. In this context, proteins associated into metabolons might exhibit interaction promiscuity, where protein surfaces near substrate entry sites can interact flexibly with multiple partners. Alternatively, as observed in the purinosome, enzymes could be tethered together with short peptide linkers to enforce spatial proximity, increase local concentrations, and minimize substrate loss into the bulk solution.^11^

By tuning interactions within synthetic enzyme complexes to operate near the critical threshold, small perturbations in interaction affinity can trigger disproportionate boosts in pathway throughput, effectively creating an ultrasensitive metabolic switch. These approaches could be further refined so that engineered metabolons form only when needed – using, say, ligand-dependent or optogenetic interaction modules – thereby boosting flux on demand without imposing constant metabolic burden.^66,67,68^

Finally, the computational framework established in this study offers a practical and broadly applicable tool for metabolic engineering, enabling performance prediction of designed enzyme assemblies using only protein-protein interaction affinities as input.

Although centered on CoQ biosynthesis, our findings suggest that weak interaction-driven clustering and enhanced substrate channeling may be general features of other multi-enzyme metabolic pathways organized in metabolons. In this perspective, the concept of metabolon – as distinct from rigid macromolecular complexes – contributes to a broader reshaping of our understanding of the role of crowding in the spatial organization of metabolic pathways within the cell.^69^ Further biophysical and computational work will be needed to test whether this holds true across pathways with diverse substrate properties.

## Methods

### Protein Expression

Ancestral tetrapod COQ proteins were expressed as previously described.^41^ Briefly, expression plasmids were cloned with synthetics DNA by Genscript as follows: COQ4, COQ6 (either full-length or C-terminally truncated) and COQ7 under a 6xHis-SUMO tag into pET-11d; COQ3, COQ4 and COQ9 under a 6xHis-SUMO tag into pET-24a(+); COQ5 under a Twinned-Strep tag fused with a PreScission cleavage site into pET-24a(+); COQ3, COQ4 (either wt and H142A-H146A), COQ5, COQ7, COQ8B, FDXR and FDX2 into a pGEX-6P-1 possessing GST-tag fused with a PreScission cleavage site upstream the cloning site. Plasmids of COQ3, COQ4 (including its double mutant), COQ6 (both constructs), COQ7 and COQ9 were transformed by heat-shock into *E. coli* BL21 cells whereas the BL21-CodonPlus-RP strain was used for all other vectors. Single colonies were inoculated in LB pre-cultures (200 rpm, 37 °C, O/N), transferred 1:100 in 1 L TB, and grown at 37 °C up to OD_600 nm_ = 0.5-0.7. Protein expression was induced with 0.1 mM IPTG. Expression temperature was 30 °C for COQ3, COQ4, COQ6, COQ7 and COQ9; 24 °C for COQ8B, FDXR and FDX2; 16 °C for COQ5. Cells were harvested by centrifugation (5,000 *g*, 10 min, 10°C) after 16 h from induction, flash frozen in liquid nitrogen and stored at −80 °C.

### Protein Purification

Ancestral tetrapod COQ proteins were purified as previously described.^41^ Briefly, cells were lysed with a high-pressure homogenizer (Emulsiflex c-3; ATA Scientific) in purification buffer (50 mM HEPES pH 7.2, 250 mM NaCl, 10% (v/v) glycerol). For soluble proteins (COQ3, COQ6 truncated, COQ9, FDXR, FDX2), the cell-free extract was obtained by centrifugation (56,000 *g*, 1 h, 4 °C). For COQ4, COQ5, COQ6 full-length, COQ7, and COQ8B, unbroken cells were removed by slow-speed centrifugation (1,200 *g*, 10 min, 4 °C) and membranes were obtained by high-speed centrifugation (56,000 *g*, 1 h, 4 °C). Proteins were extracted O/N with 1% (w/v) DDM except for COQ5 that was extracted with 1% (w/v) FOS-choline 12. Extracts were obtained by centrifugation (56,000 *g*, 1 h, 4 °C). All buffers for subsequent steps were supplemented with 0.05% (w/v) DDM.

6xHis-SUMO tagged proteins were purified in batch by elution with 300 mM imidazole in purification buffer with a step-wise gradient and desalted with a PD10 gravity column (Cytiva). Tag was cleaved by O/N incubation with 0.1 mg ml^-1^ 6xHis-SUMO protease. Tag and protease were removed by reverse affinity chromatography with an Ӓkta Pure system (Cytiva) equipped with a Ni HiTrap-HP column (Cytiva).

GST-PreScission tagged proteins were purified in batch by elution with 50 mM reduced glutathione after extensive wash with purification buffer and desalted as described above. Tag was cleaved by O/N incubation with 1 U GST-Prescission protease (Cytiva). Tag and protease were removed by reverse affinity chromatography with an Ӓkta Pure system (Cytiva) equipped with a GSTrap-HP column (Cytiva).

COQ5 was purified in batch by elution with 5 mM biotin in purification buffer after extensive wash with purification buffer and desalted as described above. Tag was cleaved by O/N incubation with 1 U GST-Prescission protease (Cytiva). Tag and protease were removed by reverse affinity chromatography with an Ӓkta Pure system (Cytiva) equipped with a GSTrap - HP column *in tandem* with a Strep-XT-Trap column (Cytiva).

Proteins were concentrated using Amicon Ultra (Merck) and their concentration was assessed using a NanoDrop ND-100 spectrophotometer (Thermo Scientific). Purity was evaluated by SDS-PAGE. Purified protein samples at about 200 μM were flash frozen in liquid nitrogen and stored at −80 °C.

Unless otherwise stated we employed C-terminally truncated COQ6 due to its convenient higher solubility and purification yields.

### Size Exclusion Chromatography coupled with Multi-Angle Light Scattering (SEC-MALS)

COQ3, COQ4, COQ5, COQ6 full-length, COQ7 and COQ9 were mixed in 50 μl of 50 mM HEPES pH 7.5, 150 mM NaCl buffer, with a final concentration of 1 mg ml^-1^ using their respectively predicted extinction coefficient at 280 nm from ProtParam (Expasy) as reference. The solution was left to stand for approximately 1 hour in ice before being spun down (16,000 *g*, 10 min at 4 °C) to remove any residual precipitate. 30 μl of 1 mg ml^-1^ COQs were injected into a Protein KW-802.5 analytical size-exclusion column (Shodex) and separated with a flow rate of 1 ml min^-1^ in phosphate buffer saline using a Prominence high-pressure liquid chromatography (HPLC) system (Shimadzu). For MW characterization, light scattering was measured with a miniDAWN multi-angle light scattering detector (Wyatt), connected to a RID-20A differential refractive index detector (Shimadzu) for quantitation of the total mass and to a SPD-20A UV detector (Shimadzu) for evaluation of the sole protein content. Chromatograms were collected and analyzed using the ASTRA7 software (Wyatt), using an estimated dn dc^-1^ value of 0.185 ml g^-1^. The calibration of the instrument was verified by injection of 10 μl of 2.5 mg l^-1^ monomeric BSA.

### Dynamic Light Scattering

Dynamic Light Scattering measurements were performed with a Prometheus Panta NT.48 Series instrument (NanoTemper) set at 25 °C and High Sensitivity Capillaries (NanoTemper). COQ3, 4, 5, 6, 7, 9 have been incubated in phosphate buffered saline (PBS) for 15 min at room temperature at a final concentration of 1 mg ml^-1^. The residual detergent was removed using a PD SpinTrap G-25 column (Cytiva) according to manufacturer’s instructions, and aggregates by centrifugation (20,000 *g*, 10 min, 4 °C). Each measurement consisted of an average of 10 scans of 5,000 ms collected with laser power set at 100% with PR.Panta Control software (NanoTemper). COQ metabolon was measured in 4 independed replicates, while single proteins and buffer controls in triplicate. Data fitting to autocorrelation function was carried out using PR.Panta Analysis software (NanoTemper). Data sets showing a polydispersity index (PDI) > 0.2 were fitted with a size distribution model, while monodispersed ones (PDI < 0.2) with a cumulant analysis model.

### Micro-Scale Thermophoresis

COQ3, COQ5, COQ7 and COQ9 were labeled on their primary amino groups with 2^nd^ generation RED-NHS dye (NanoTemper) following the provider’s protocol. Briefly, 10 μM protein was incubated at RT with 300 μM dye in the kit labeling buffer. Excess fluorophore was removed by desalting with a B-column (NanoTemper) pre-equilibrated in assay buffer (50 mM Tris-HCl pH 8.5 at 4°C, 100 mM NaCl, 5% (v/v) glycerol, 0.05% (v/v) Tween-20, 0.1% (w/v) Pluronic acid). The protein concentration and degree of labeling (DOL) was estimated based on the protein’s absorbance at 280 nm and the fluorophore’s at 650 nm. Labeled proteins with a DOL of at least 0.5 were employed in binding assays. Protein-protein binding affinity experiments were performed with 40 nM final labeled protein titrated with a serial 1:2 dilution of the unlabelled ligand protein in the low μM and nM range. Experiments were conducted with Premium MST capillaries (NanoTemper). All experiments were performed at least in triplicate.

COQ3, COQ4, COQ6, COQ7 and COQ9 baring 6xHis-SUMO tag were further purified by size exclusion chromatography with an Ӓkta Pure system (Cytiva) equipped with a Superdex 200 Increase 5/150 GL column (Cytiva) equilibrated in phosphate buffer saline with 0.05% (v/v) Tween-20 (PBS-T). His tagged proteins were non-covalently labeled with 2^nd^ generation RED-tris-NTA (NanoTemper). The affinity of each protein for the dye was determined by titrating the dye with serial 1:2 dilutions of each protein (2 μM - 61 pM range) in PBS-T. His-tagged COQs with an affinity with the dye of at least 10 nM were employed as target in subsequent binding assays. Binding affinity experiments were performed in PBS-T with 50 nM His-tagged target protein, pre-incubated with 5 μM dye for 30 min at RT, titrated with serial 1:2 dilutions of the unlabelled ligand protein in the low μM to nM range.

### Analytical Size Exclusion Chromatography

The COQ metabolon (COQ3-7 and COQ9) was reconstituted in its intact and proten-deprived forms by mixing proteins at 1 mg ml^-1^ final concentration in 50 mM HEPES pH 7.5, 150 mM NaCl buffer, at a final concentration of 1 mg ml^-1^ (50 μl final). The mixes were incubated for 1 hour in ice before being spun down (16,000 *g*, 10 min at 4°C) to remove any residual precipitate. Samples were loaded on an ÄktaGo system (Cytiva) equipped with a Superdex 200 Increase 5/150 GL column (Cytiva) pre-equilibrated in phosphate buffer saline. Chromatography was carried out at a flow rate of 0.4 ml min^-1^ and 200 μl fractions were collected at 1.4, 1.6 and 1.8 ml of each run. All fractions (15 μl) were analyzed by SDS-PAGE and submitted to bottom-up proteomics studies.

### Bottom-up Proteomics

Bottom-up proteomics was performed for various COQ assemblies separated by analytical SEC. Each fraction was diluted with a buffer to achieve 100 mM Tris, pH 8.5, 10 mM TCEP, 40 mM CAA, 1% SCD (all final concentrations), and incubated for 30 min at RT. After, the sample was 10x diluted with 50 mM ammonium bicarbonate, pH 8.5. Sample digestion was performed with trypsin 1:20 (w/w) and LysC 1:50 (w/w) at 37 °C for 16 h. The digestion was quenched by TFA (0.5 % (v/v) final concentration), and samples were desalted on an Oasis HLB µElution Plate (Waters Corp) using a vendor-provided protocol. The solvent was fully evaporated, and samples dissolved in 2% (v/v) FA prior to the liquid chromatography-mass spectrometry (LC-MS) analysis.

The LC-MS analysis was performed in triplicate on an UltiMate 3000 UHPLC system coupled to an Orbitrap Exploris 480 (Thermo Fisher Scientific Inc). The peptides were analyzed in 90 min gradient. First, the peptides were trapped on an Acclaim Pepmap 100 C18 (5 mm × 0.3 mm, 5 μm, Thermo Fisher Scientific Inc) for 1 min at a flow rate of 30 μl min^-1^, solvent A (0.1% FA v/v in H2O). The trapping was followed by separation on an analytical column packed in-house (Reprosil 2.4 μm, 75 μm × 50 cm) at a flow rate of 0.3 μl min^-1^ using a gradient starting from 9% solvent B (80% acetonitrile (ACN) v/v, 0.1% FA v/v) at 0–1 min, 13% B at 2 min, 44% B at 67 min, 55% B 72 min, 99% B 75 min, and equilibrated for the next run in 9% B at 80–90 min. MS1 scans were collected with the following parameters: resolution = 60000, mass range = 375-1600 Th, standard AGC target, automatic maximum injection time, cycle time = 1 s. MS2 scans were collected in data-dependent mode for precursors with charge state 2-6, with the following parameters: isolation window = 1.4 Th, resolution = 15000, mass range = 120-2000 Th, HCD collision energy = 28 %. Precursors that were once selected for fragmentation were excluded from MS2 for 14 seconds.

The resulting data were searched in FragPipe 22.0 against reviewed human FASTA (UniProt, accessed 13-10-2023 with common contaminants), where the COQ3-7, 8B, and 9 were replaced by the recombinant ancestral protein sequences.^41,70^ The peptide mass and length were set to be between 500-5000 Da and 7-50 amino acids, respectively. Precursor and fragment tolerances were set to 10 ppm and 20 ppm, respectively. Carbamidomethylation [C] was set as fixed modification, and oxidation [M] was allowed as variable modification. Results were filtered for 1% FDR and quantified using label free quantification (MaxLFQ).

### Cross-linking Mass-Spectrometry

XL-MS experiments were performed both in-solution, and in-gel in an experimental triplicate. The in-solution crosslinking was first optimized (Fig. S14). Based on the optimization, 0.5 μg μl^-1^ (∼3 μM) COQ sample in 50 mM Hepes pH 7.5 was cross-linked with 0.5 mM DSS or 2.5 mM DMTMM for 1 h at room temperature (RT). The reaction was quenched by addition of 1 M Tris, pH 8.5 to achieve 25 μM final concentration. The samples were processed as described above for bottom-up proteomics with the only difference in enzyme:protein ratios adjusted to 1:20 trypsin (w/w), 1:50 LysC (w/w).

In gel cross-linking was performed for different assemblies of COQ proteins separated by Blue native-PAGE (Fig. S15), as previously described.^44^ Briefly, the bands of interest were excised, washed with cold PBS pH 7.5 and submerged in 50 mM Hepes pH 7.5 containing either 1.5 mM DSS or 5 mM DMTMM for 1 h at RT. The reaction was quenched by addition of 1 M Tris pH 8 to achieve 25 μM final concentration. The bands were washed with MQ H_2_O and dehydrated by 100% ACN. After, the samples were reduced by 6.5 mM dithiothreitol in 50 mM ammonium bicarbonate pH 8.5 (1 h, 60 °C). After reduction, the samples were dehydrated by 100% ACN and alkylated by 55 mM iodoacetamide in 50 mM ammonium bicarbonate pH = 8.5 (30 min, RT, in dark). The alkylation was followed by washing and dehydrating the pieces two times with 50 mM ammonium bicarbonate, pH = 8.5. Importantly, each dehydration step involved submerging the gel pieces in 100% ACN (15 min, RT). The dehydrated gel pieces were submerged in trypsin solution (3 ng μl^-1^ in 50 mM ammonium acetate, pH 8.5) and incubated (90 min, on ice). After, 25 μl of 50 mM ammonium bicarbonate pH 8.5 was added to each sample and incubated (16 h, 37 °C). Next, the supernatant with digested peptides was collected, and gel pieces were dehydrated with 100% ACN. The supernatants were combined, and the solvent was fully evaporated. Dry peptides were dissolved in 2% FA prior to the LC-MS/MS analysis.

Peptides from in-gel and in-solution cross-linking were processed by LC-MS as described above for bottom-up proteomics. The XL-MS method included several adjustments: charge states selected for MS2 = 3-8, MS2 scan range = 200-2000 m/z, MS2 resolution = 30000, HCD collision energies = 24, 28, 33 %, and precursors that were once selected for fragmentation were excluded from MS2 for 30 seconds.

The raw data were searched using pLink 2.0 against a database containing sequences of ancestral COQ3:7 and COQ9 sequences and common contaminants.^71^ The linker mass for DSS (K-K) and DMTMM (E|D–K) was set to be 158.004 Da and −18.011, respectively. N-terminus linkage was allowed. Trypsin was set as the enzyme, cleaving at the C-terminal of K and R with up to 3 allowed missed cleavages. The peptide mass and length were set to be between 600-6000 Da and 6-60 amino acids, respectively. Precursor and fragment mass tolerances were set to be 20 ppm. Carbamidomethylation [C] was set as fixed modification, and oxidation [M] as variable modification. The results were filtered for 1% FDR and scores <10e-4. A cross-linked peptide pair was considered, only if detected in at least 2 out of 3 experimental replicates.

### Protein PEGylation

Purified COQ proteins were PEGylated on surface primary amino groups using O-methyl-O’-succinyl polyethylene glycol 5,000 N-succinimidyl ester (methoxyPEG-NHS). Protein concentration was adjusted to 10 mg ml^-1^ in purification buffer supplemented with 100 mM sodium bicarbonate. 100 μl of protein were incubated with 10 μl of 20 mg ml^-1^ methoxyPEG-NHS for an hour at RT. The reaction was quenched by adding 10 μl of 1 M Tris-HCl pH 8 at 4 °C. The excess of reactant was removed by size exclusion chromatography with an Ӓkta Pure system (Cytiva) equipped with a Superdex 200 Increase 5/150 GL column (Cytiva) equilibrated in 50 mM Tris-HCl pH 8 at 4 °C.

### Pull-down Assay

His-SUMO tagged COQ6 was used as bait, while both non-modified and PEGylated tag-less COQ3 served as prey in independent experiments. Both proteins were mixed at 10 μM in 1 ml volume of purification buffer, incubated for 30 min in ice and loaded into a gravity column containing 1 ml Ni Sepharose excel resin (Cytiva) pre-equilibrated in purification buffer. The resin was washed with 5 column columns of purification buffer and elution was performed with the same volume of purification buffer supplemented with 300 mM imidazole. Flow-through, wash, and elution samples were concentrated back to 1 ml with Amicon Ultra 15 10 kDa cut-off. SDS-PAGE analysis was performed to analyze the content of each chromatographic step. The experiment was performed in triplicate (Supplementary Fig. 20).

### Blue Native Poly-Acrylamide Gel Electrophoresis

Proteins were pre-incubated on ice in 50 mM Tris-HCl pH 8.0 (4°C) at a final concentration of 1 μM for one hour. Samples loaded with the NativePAGE Sample Buffer (Invitrogen) on a 1 mm NativePAGE Bis-Tris mini protein 4-16% precast protein gel (Invitrogen). Anode buffer consisted of 50 mM Bis-Tris and 50 mM Tricine pH 6.8; cathode buffer consisted of anode buffer supplemented with 0.02% (w/v) Coomassie G-250. Electrophoresis run was carried out at 4 °C at 150 V for 60 min and 250 V for 90 min. Fixing, Coomassie R-250 staining and destaining were performed according to the manufacturer instructions.

### Peptide Mapping

Excised bands were de-stained in 100 mM ammonium bicarbonate pH 7.8, 50% (v/v) acetonitrile and dehydrated with pure acetonitrile. Gel fragments were re-hydrated with 20 ng μl^-1^ sequencing grade trypsin (Promega) in 100 mM ammonium bicarbonate pH 7.8 and incubated overnight at 37 °C at 200 rpm. Peptides were extracted twice with 50% (v/v) acetonitrile, 5% (v/v) formic acid in water by vortexing for 15 min. Pooled extracts were dried and resuspended in 50 μl UHPLC grade water with 0.1% (v/v) formic acid.

Samples were analyzed by ESI-qTOF on a X500B system (SCIEX) equipped with the Twinned Sprayer ESI probe coupled to an ExionLC system (SCIEX) controlled by SCIEX OS software 3.4. Injection volume was 45 μL. A Jupiter Proteo 90 Å column (150 mm length x 2 mm diameter, 4 μm particle size; Phenomenex) was employed for chromatographic separation. Mobile phase A consisted of water, B of acetonitrile, both supplemented with 0.1% (v/v) formic acid. Flow rate was 0.2 ml min^-1^. Gradient elution was performed with a linear 2-15% B gradient in 15 min. MS detection was performed as follows: curtain gas 30 psi, ion source gas 1 140 psi, ion source gas 2 45 psi, temperature 350 °C. Full-scan and fragmentation spectra were generated using information dependent acquisition (IDA) in positive mode with spray voltage +4,500 V, declustering potential of +10 V and collision energy of +10 V. Mass calibration was performed with the ESI positive calibration solution (SCIEX) prior to experiments. Mass spectra were processed with Peaks studio 4.5 and resulting peptides were searched against an ad hoc database of ancestral COQ proteins’ sequences.

### Small-scale Reactions

Small-scale *in vitro* reconstructions of the COQ metabolon’s pathway were performed similarly to previously reported methodology.^41^ The reaction mix contained 1 μM COQ3, COQ4, COQ5, COQ6, COQ7, COQ8B and COQ9, 2 μM FDXR and FDX2, 250 μM FAD, 1 mM S-adenosylmethionine, 150 μM MgCl_2_, 25 μM ZnCl_2_, 1 mM substrate, NAD(P)H and ATP regeneration systems. The NAD(P)H regeneration system consisted of 300 μM NADP+ and NAD+, 1.2 U glucose dehydrogenase and 1.2 mM glucose. The ATP regeneration system consisted of 1 mM ADP, 4 U pyruvate kinase, 5 mM phosphate and 5 mM phosphoenolpyruvate. Control reactions were performed with all non-modified COQ proteins. Samples were prepared by using one single PEGylated COQ protein at a time, with COQ4 H142A-H146A, and without FDXR and FDX2. The activity of each PEGylated COQ protein was assayed by incubation with their cofactor(s) and substrate(s) at the previously mentioned concentrations. Reactions were incubated at 25 °C 200 rpm O/N in a bench-top shaking block in low-light conditions and quenched by adding pure acetonitrile 1:3. Proteins were removed by centrifugation at 20,000 *g* 10 min and 1 μM sorbicillin was added as an internal standard. All samples were diluted 1:10 in purification buffer and acetonitrile 2:1 containing 1 mM Sorbicillin prior to injection in UHPLC/HRMS.

UHPLC/HRMS analysis were carried out on the previously described system similarly as previously described.^41^ Injection volume was 10 μL. A Kinetex EVO C18 column (100 mm length x 2.1 mm diameter, 2.6 μm particle size; Phenomenex) was employed for chromatographic separation. Mobile phase A consisted of water, B of acetonitrile, both supplemented with 0.1% (v/v) formic acid. Flow rate was 0.2 ml min^-1^. Gradient elution was performed as follows: 2% B at 0.0-0.1 min, 2-66% B at 0.1-32.0 min, 66-2% B at 32.0-35.0 min. MS detection was performed as follows: curtain gas 30 psi, ion source gas 1 45 psi, ion source gas 2 55 psi, temperature 450 °C. The full-scan range of *m/z* 50-1,000 was monitored in positive mode with spray voltage +5,500 V, declustering potential of +50 V and collision energy of +10 V. Mass calibration was performed with the ESI positive calibration solution (SCIEX) prior to experiments. The content of **1**, **2**, **3**, **4a**, **4b**, **5_ox_** and CoQ_1_ in the reactions was quantitated in the reaction using Sciex OS 3.4 Analytics software. Standard solutions were prepared in the 3-100 μM range in purification buffer supplemented with sorbicillin and diluted 1:3 in acetonitrile supplemented with 1 μM sorbicillin and injected in duplicate. Calibration curves were generated utilizing the extracted ion chromatogram (XIC) of the [M+H]^+^ *m/z* of each analyte and sorbicillin as an internal standard, R^2^ values were at least 0.98 (Fig. S16). The mass error threshold for intermediates identification was set at ± 5 ppm. Intermediates were quantified based on the XIC of *m/z* corresponding to the [M+H]^+^ of each analyte extracted from all samples.

### Steady-state Kinetics

The activity of PEGylated COQ6 and COQ7 was checked following published protocols.^41^ Briefly, final protein concentration was 5 μM in 150 μl of purification buffer. For COQ6 activity assay 50 μM FAD, equimolar FDXR and FDX2, 200 μM **1** were pre-incubated and reaction was started by adding 50 μM NADPH. For COQ7 activity assay equimolar COQ7/COQ9 and 700 μM **5_ox_** were pre-incubated and the reaction was started by adding 500 μM NADH. Activity was measured by monitoring spectrophotometrically NAD(P)H oxidation (ε_340 nm_ = 6.22 mM^-1^ cm^-1^) in 10.00 mm quartz cuvettes (Hellma) and a Cary 100 spectrophotometer (Agilent) equipped with a thermo-stated cell holder (T = 25 °C). Experiments were performed in duplicate.

### NMR

Sorbicillin was dissolved in deuterated methanol (MeOD) and ^1^H spectra were acquired with 100 scans on a Bruker 400 MHz Avance III instrument and analysed using TopSpin 4.3.0 (Supplementary Fig. 17).

### Simulations

#### Simulation Environment

All Langevin dynamics simulations were run in LAMMPS.^72^ We modeled COQ3-7 as hard spheres of radius 10 Å with an adhesive interaction site of radius 3.5 Å that represents the active site of the enzyme. COQ9 and crowder were modelled as hard spheres of radius 10 Å without an active site. Substrate molecules, also modeled as hard spheres of radius 1 Å, could diffuse throughout the 800A cubic simulation box.

#### Initial Conditions

Each simulation began with 50 particles for COQ3-7 and COQ9, 1,000 particles of compound **1**, and 700 particles of crowder. The masses of COQ3-7, COQ9, and crowder were set to 30,000, while the active sites had a mass of 1,500 and the substrates/intermediates/products had a mass of 800. Each active site patch was made complementary to a specific substrate particle type or types, where for example compounds **1** and **4a** were native to COQ6 and compound **2** was native to COQ3. Enzyme catalyzed reactions were enabled using bond/create/random and bond/break functionalities in LAMMPS.^72,73^ When a native substrate-patch pair binds by coming into contact within 5.5 Å, a reaction occurs under a certain probability where the type of the substrate particle was changed into the corresponding product’s type. In order to model reaction-limited (not diffusion-limited) enzymes, the default probability was set to be p_base_ = 0.00001 (i.e. *k_cat,base_*).^44^ This value was allowed to vary in our simulation, where for instance 5*k_cat,base_* would correspond to simulations using a reaction probability of 0.00005.

#### Solvent and Interaction Modeling

A Langevin thermostat was applied to all particles to ensure an NVT ensemble at 310 K and model interactions with implicit solvent. The total force on each particle can be decomposed into three parts: 1) the conservative force *F*_*c*_ arising from inter-particle potential energy functions, 2) the frictional drag *F*_*f*_, also known as viscous damping force, and 3) force *F*_*r*_ due to random collisions with solvent particles.

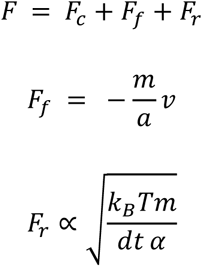

The damping factor *α*, which is in time units, sets how quickly the temperature gets relaxed in the simulation. *α* was set to 2000 fs for COQ3-7, COQ9, and crowder and to 800 fs for all substrates/intermediates/products. The smaller the alpha is, the higher the viscosity of the modeled solvent becomes. The values were selected to model the viscosity of water, which is 10^-3^ Pa s at 310 K, using the below relation.

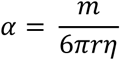

All pairwise interactions between particles were modeled using Lennard Jones potentials with a cut off distance that is roughly 2.5 times the minimum distance, with the exception of native active site patch and substrate interactions that used Morse potentials. The COQ enzymes (excluding active site patches) could interact among themselves with an interaction strength of ε_enz-enz_ that were calculated in a way that reflected the experimentally measured K_d_ constants. We converted this by using the Arrhenius equation k_off_ = A * e^(-ε/kT)^, where k_off_ is the dissociation rate constant, A is the Arrhenius factor that is assumed to be 1, and ε is the interaction strength. The protein-protein pair specific k_off_ is estimated with k_off_ = K_d_ * k_on_ where the theoretical protein-protein k_on_ is assumed to be 10^5^.^74^ Thus, the interaction strength is estimated based on ε_enz-enz_ = -kT * ln(K_d_ * k_on_). The self-interaction strength of COQ3-7, and COQ9 were all set as 1kT. Ligands interacted with both the enzyme active site and the nonspecific protein surface. The interaction strength ε_patch-lig_ between a native/cognate active site and ligand pair was set as 4kT, while the nonnative/noncognate interaction was set as 2kT. All interactions between an active site patch and the final product representing coenzyme Q only included a repulsive part of the potential without the attractive portion. The protein surface of COQ3-7 and COQ9 (excluding the active site) interacted with all ligand particles with a strength of ε_enz-lig_ = 2kT, except with coenzyme Q that only had a repulsive portion. The crowder interacted with all ligands with a strength of ε_crowd-lig_ = 1/3kT. All other types of pairwise interactions not mentioned here explicitly only included a repulsive portion. Details of all parameters could be found in the code we released.

#### Data Collection

The integration timestep for simulations was set to 25 fs. To ensure proper initialization, an equilibration phase of 625 ns was conducted with enzyme-catalyzed reactions turned off, allowing for the formation of enzyme clusters. Initial particle coordinates were generated using a Python script to ensure non-overlapping particles. Following equilibration, the production run commenced with enzyme-catalyzed reactions activated. Data was collected every 10,000 steps during the production run, recording the cumulative production count of each ligand type and reaction data (timestep, enzyme particle id and substrate particle id). This data was subsequently post-processed to calculate the number of each ligand type present in the simulation box over time. The simulation was repeated multiple times to ensure reliability and reproducibility of the results.

#### PEGylation simulations

PEGylation was modelled by turning off all enzyme-enzyme interactions involving the PEGylated protein. For instance, simulations with PEGylated COQ6 set all ε_COQ6-enz_ to 0.01 kcal/mol, where enz stands for all other COQs. Any other interaction strengths were unchanged. The production run for PEGylation simulations lasted 7.5 us, and the enzyme activity value (represented by the reaction probability) was set at 10 *k_cat,base_*. Each simulation condition was replicated 3 times with different seeds.

#### No interaction simulations

The production run for simulations with all ε_enz-enz_ set as 0.01 to only include repulsive interactions lasted 7.5 us. The enzyme efficiency value was set at 10 *k_cat,base_*. Each simulation condition was replicated 3 times with different seeds.

#### Randomization simulations

Randomization of the experimentally measured protein-protein interaction network was done using a python script. ε_enz-enz_ values were randomly shuffled except the ε_COQ7-COQ9_ = 4.45 kcal/mol, because COQ9 is known to be crucial for the functional efficiency of COQ7 by enhancing its catalytic activity. The production run lasted 15 us and the reaction probability was varied across the following multiples of *k_cat,base_*: 0.5, 1, 2, 5, 10. For each set of parameters, the simulations with a randomized interactions network were replicated 12 times with different seeds. The simulations with the experimentally measured interaction network were replicated 6 times.

#### Ablation simulations

Protein-protein interactions were separately ablated by setting their ε to 0.01 kcal/mol and restricting the potential to its repulsive component. The production run lasted 15 us, and the reaction probability was set at 10*k_cat,base_*. The two simulation conditions were replicated 3 times each with different seeds.

#### Channeling postprocessing

For all ligand particles that become **3**, **4a**, **4b**, **5_ox_**, **6_ox_**, or coenzyme Q, that underwent at least two reactions, let *P* denote a substrate that undergoes a series of *n* consecutive reactions within the pathway. For each reaction step *i* ∈ {1, 2,… *n*-1}, let *τ_i_*(*P*) be the simulation timeframe at which the *i-th* reaction occurs for particle P. We can define the dwelling time as:

For example, if the 7 reactions that converted particle *P* from compound **1** to coenzyme Q happened at timeframes t, t+1, t+2, t+4, t+10, t+11, and t+14, the values 1, 1, 2, 6, 1, and 3 would be recorded.

#### Enzyme clustering postprocessing

Using a custom VMD tcl script, enzyme clusters (amongst COQ3-7 and COQ9) were computed at each simulation timeframe and their compositions based on particle IDs were recorded. If the distance between two enzymes was greater than 25 Å, they were deemed to be in different clusters. A group of enzymes computed this way was only considered to be a cluster if it had more than 4 members. These enzyme clusters were further filtered based on completeness, where a cluster was considered to be complete if it included at least one copy of COQ3-7.

#### Bulk reaction to cluster reaction ratio postprocessing

To quantitatively analyze reaction localization within the simulation environment, we defined a binary classification for each catalytic event *R_i_.* Let *E_i_* denote the enzyme catalyzing reaction *R_i_*, and *C(E_i_, t_i_)* represent the cluster containing enzyme *E_i_ at time t_i_ when the reaction occurs.* We can classify each reaction according to:

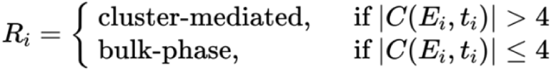

Where |*C(E_i_, t_i_)|* denotes the number of enzyme members in the cluster. We calculate the bulk-to-cluster reaction ratio for the entire simulation as

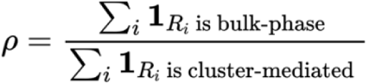

where **1** is the indicator function

#### Yield Ratio Calculation

To calculate the ratio (r) between the randomized and experimental conditions, we used the final and mean concentrations of the Coenzyme Q product generated during simulations conducted over 12 randomized runs and 6 wild-type runs at different *k_cat_* values. We can define the ratio as

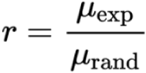

The standard error of the ratio σ_r_ was calculated using error propagation theory:

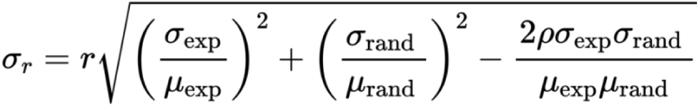

Where s_exp_ and s_rand_ are the standard deviations of experimental and randomized yields, respectively and ρ is the Pearson correlation coefficient. This appoach accounts for both the individual variances and their covariance. In our results, we present the mean advantage ratio, calculated by averaging the yield ratios across all sampled timepoints throughout the simulation.

## Data availability

The experimental data generated in this study are provided in the supplementary information and source data files. All other data are available from the authors upon reasonable request. The ancestral sequences have been deposited to the Genbank database under accession codes OQ859710, OQ859711, OQ859712, OQ859713, OQ859714, OQ859716, OQ859717, OQ859718, OQ859719. The bottom-up proteomics and XL-MS data have been deposited to the ProteomeXchange Consortium via the PRIDE partner repository with the dataset identified PXD062263.^75^ Source data are provided with this paper.

## Code availability

The original code developed for this work is deposited in https://github.com/jiojeongharvard/CoQ_metabolon and is available as the date of the publication.

## Acknowledgments

This research was funded by the ERC Advanced Grant, MetaQ (no. 101094471), and by the Associazione Italiana per la Ricerca sul Cancro (AIRC) Investigator Grant (no. 28754) to A.M. The authors thank Dr. Mannucci B. (*Centro Grandi Strumenti* Core Facility, University of Pavia) for the technical support in analytical chemistry methods development and Professor Forneris F. (Department of Biology and Biotechnology, University of Pavia) for the technical support in SEC-MALS data acquisition. This research was also funded by NIH R35GM139571 to E.I.S. The content is solely the responsibility of the authors and does not necessarily represent the official views of the National Institutes of Health.

## Author information

### Constitutions

Conceptualization: all authors; Methodology: D.W., A.G., J.J., C.R.N., J.L., T.K., A.M. E.I.S.; Investigation: D.W., A.G., J.J., C.R.N., J.L., T.K., D.C., M.M.; Writing the original draft: D.W., A.G., J.J., C.R.N.; Review & Editing the original draft: T.K., A.J.R.H., A.M., E.I.S.; Data Visualization: D.W., A.G., J.J., C.R.N., T.K.; Supervision: A.J.R.H., A.M., E.I.S.; Funding Acquisition: A.M., E.I.S.

### Corresponding authors

Correspondence to Andrea Mattevi and Eugene I. Shakhnovich.

