## Supplementary Information for "Complete Enzyme Clustering Enhances Coenzyme Q Biosynthesis via Substrate Channeling"

### Table of Contents


**Supplementary Figures** Pages 2-20

**SF1.** Individual COQ proteins show a monodisperse profile in DLS 2
**SF2.** In-gel XL-MS. 3
**SF3.** Binary protein-protein affinity measurements using micro-scale thermophoresis and amines labelling strategy 4-5
**SF4.** Protein-protein affinity measurements for COQ protein pairs using micro-scale thermophoresis and His-tag labelling strategy 6-7
**SF5.** Interaction network determines metabolon fine structure. 8
**SF6.** Fine structure of enzyme cluster has no effect on coenzyme Q yield. 9
**SF7.** Correlation between coenzyme Q yield and the proportion of enzymes in complete clusters (containing at least one copy of COQ3-7) at increasing activity values. 9
**SF8.** Progression of average number of complete clusters over time. 10
**SF9.** Reaction scheme of PEGylation of primary amino groups. 11
**SF10.** AlphaFold-3 predicted structures of COQs shown as surfaces. Surface lysine residues are colored in cyan. 11
**SF11.** Pull-down assay demonstrates that PEGylation of COQ3 impairs its binding with COQ6. 12
**SF12.** PEGylated COQs retain their enzymatic activities. 13
**SF13.** Analytical size-exclusion chromatography followed by bottom-up proteomics of the COQ metabolon depleted of individual proteins 14
**SF14.** Uncropped reducing SDS-PAGE gel of cross-linking reaction optimization for COQ3-7, 9 sample 15
**SF15.** Blue native-PAGE of COQ3-7, 9 sample 16
**SF16.** Automated peak integration and calibration curves of all analytes used for LC/MS quantitation experiments 17
**SF17.** $^{1}$H NMR spectra of the internal standard Sorbicillin. 18
**SF18.** Uncropped Blue Native PAGE gels. Each lane reported in the main figures is marked by a red arrow 19
**SF19.** Uncropped SDS-PAGE analysis of fraction from chromatograms of Fig. 2d and Fig. 6d. 19
**SF20.** Uncropped SDS-PAGE analysis of the COQ6-COQ3 pull-down assay 20

**Supplementary Tables** Page 21

**ST1.** Peptide mapping analysis of the Blue Native PAGE gel from Fig. 6b. 21
**ST2.** Peptide mapping analysis of the Blue Native PAGE gel from Fig. 7b. 21

### Supplementary Figures

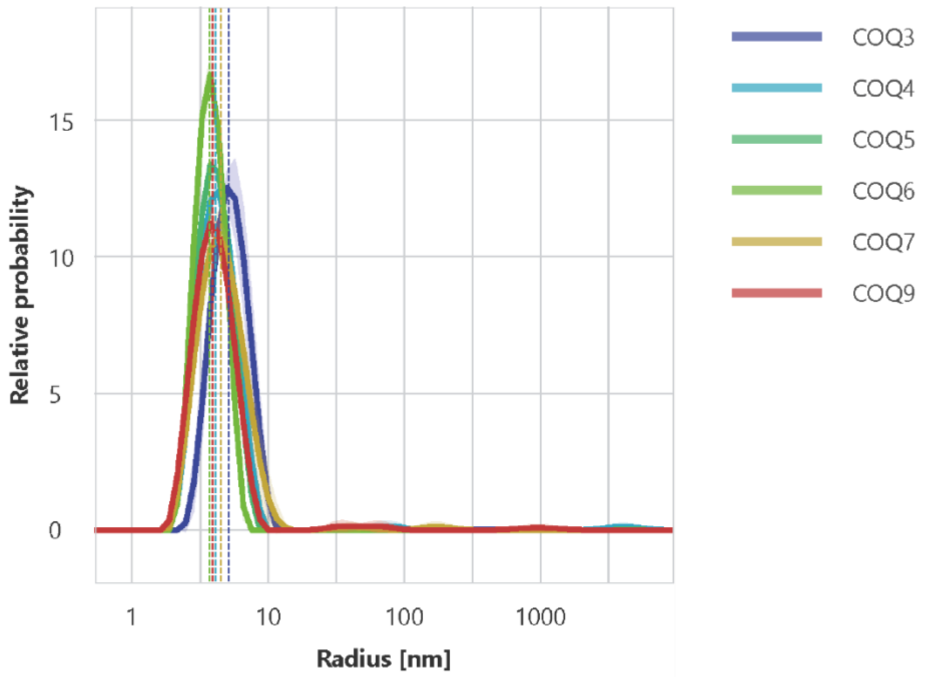


**Supplementary Figure 1. Individual COQ proteins show a monodisperse profile in DLS.** Individual COQs control runs. Data are shown as average (line) and SD (shaded area) of n=3 independent measurements, each resulting from the averaging on n=10 individual measurements. Data are fitted with a cumulant fit model resulting in a hydrodynamic radius value of 5.01±0.33 nm (COQ3), 4.09±0.07 nm (COQ4), 3.86±0.02 nm (COQ5), 3.65±0.01 nm (COQ6), 4.27±0.12 nm (COQ7) and 3.85±0.04 nm (COQ9).

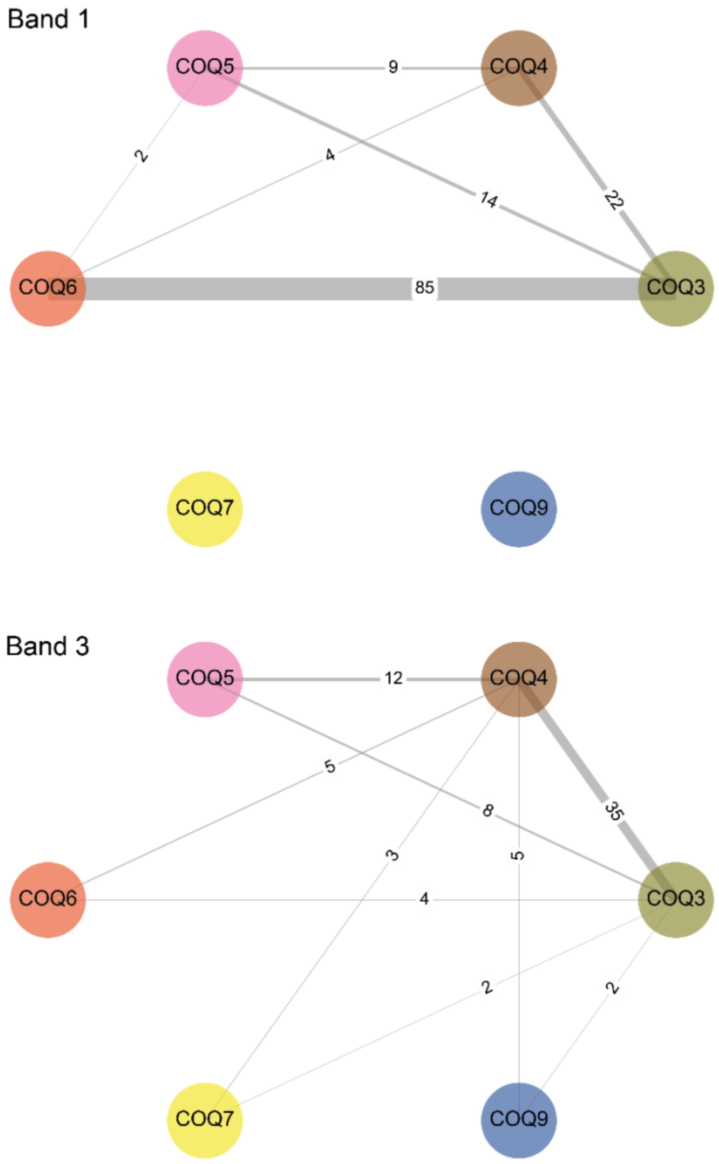


**Supplementary Figure 2. In-gel XL-MS.** Network plots showing cross-link matched spectra (CSMs) detected by in-gel cross-linking mass spectrometry of COQ metabolon of gel bands 1 (top) and 3 (bottom) separated by blue native PAGE (Supplementary Fig. 15). Edges of the network are labelled with the number of CSMs found in the analysis. The corresponding data for the gel band 2 are provided in the main Figure 2c.

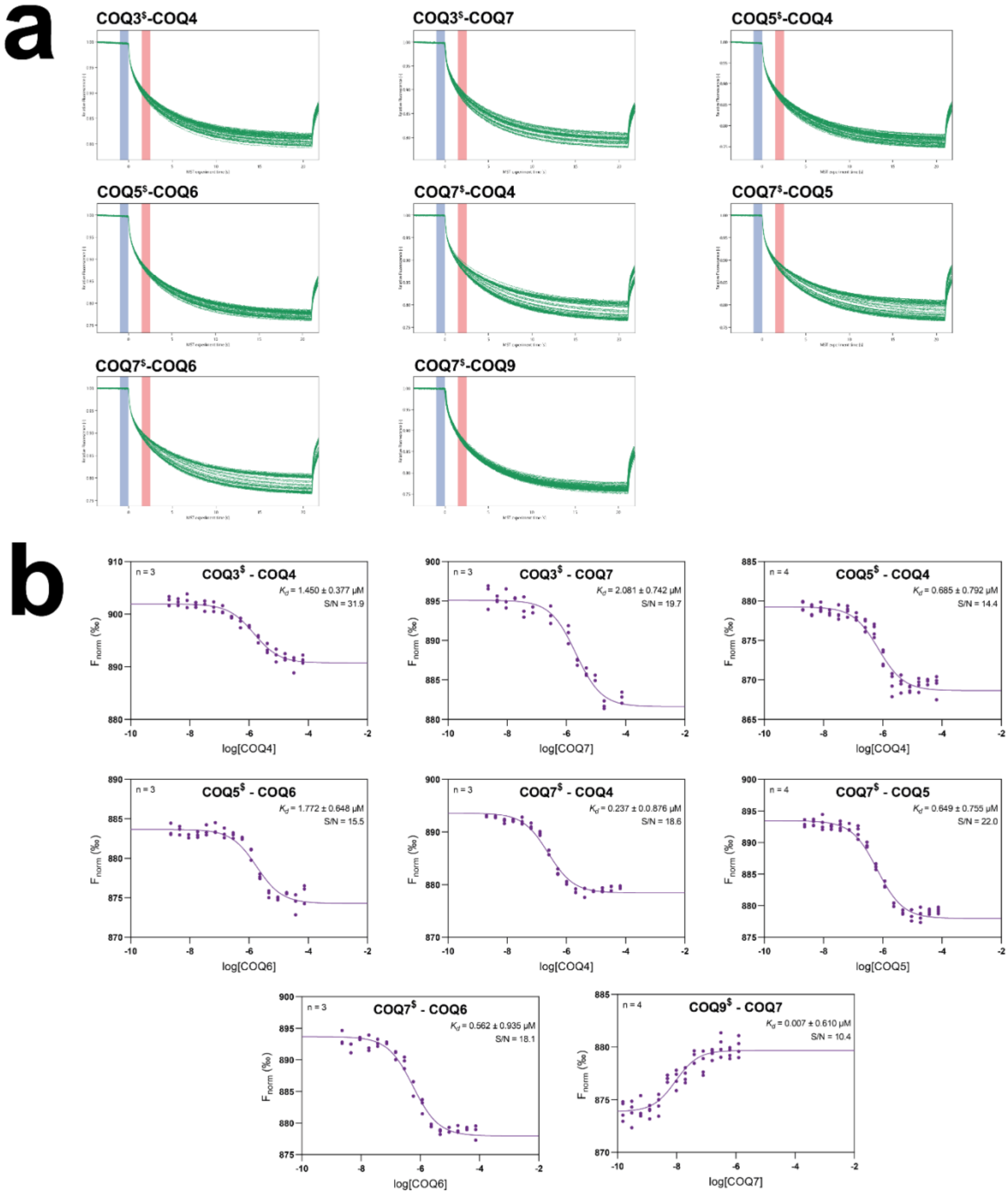

a
COQ3$-COQ4
COQ3$-COQ7
COQ5$-COQ4
COQ5$-COQ6
COQ7$-COQ4
COQ7$-COQ5
COQ7$-COQ6
COQ7$-COQ9
b
n = 3
COQ3$ - COQ4
Kd = 1.450 ± 0.377 μM
S/N = 31.9
log[COQ4]
n = 3
COQ3$ - COQ7
Kd = 2.081 ± 0.742 μM
S/N = 19.7
log[COQ7]
n = 4
COQ5$ - COQ4
Kd = 0.685 ± 0.792 μM
S/N = 14.4
log[COQ4]
n = 3
COQ5$ - COQ6
Kd = 1.772 ± 0.648 μM
S/N = 15.5
log[COQ6]
n = 3
COQ7$ - COQ4
Kd = 0.237 ± 0.0.876 μM
S/N = 18.6
log[COQ4]
n = 4
COQ7$ - COQ5
Kd = 0.649 ± 0.755 μM
S/N = 22.0
log[COQ5]
n = 3
COQ7$ - COQ6
Kd = 0.562 ± 0.935 μM
S/N = 18.1
log[COQ6]
n = 4
COQ9$ - COQ7
Kd = 0.007 ± 0.610 μM
S/N = 10.4
log[COQ7]
Fnorm (‰)

**c**

**Primary Amines Labelling**

| PROTEIN A$ | PROTEIN B | $K_d$ (µM) |
|---|---|---|
| COQ9 | COQ7 | 0.007 ± 0.610 |
| COQ7 | COQ4 | 0.237 ± 0.876 |
| COQ7 | COQ6 | 0.562 ± 0.935 |
| COQ7 | COQ5 | 0.649 ± 0.755 |
| COQ5 | COQ4 | 0.685 ± 0.792 |
| COQ3 | COQ4 | 1.450 ± 0.377 |
| COQ5 | COQ6 | 1.772 ± 0.648 |
| COQ3 | COQ7 | 2.081 ± 0.742 |

$surface Lys residues labelled with RED-NHS 2nd generation (NanoTemper)

**Supplementary Figure 3. Binary protein-protein affinity measurements using micro-scale thermophoresis and amines labelling strategy**. The Lys fluorescently labelled COQ protein is marked with $. **a.** Micro-scale thermophoresis traces; the time used for plotting dose-response curves is highlighted in red, the reference signal used for normalization in blue. **b.** Dose-response curves. The labelled protein is marked with $; dissociation constants and signal to noise ratio (S/N) are reported. The number of independent replicates is reported for each pairwise interaction. **c.** Dissociation constants ($K_d$s) are ranked from tightest to looser interactions. Data are reported as mean and SD of n≥3 (see panel b) independent experiments. The values measured using His-tag labelling are listed in Fig. 2d.

**a**

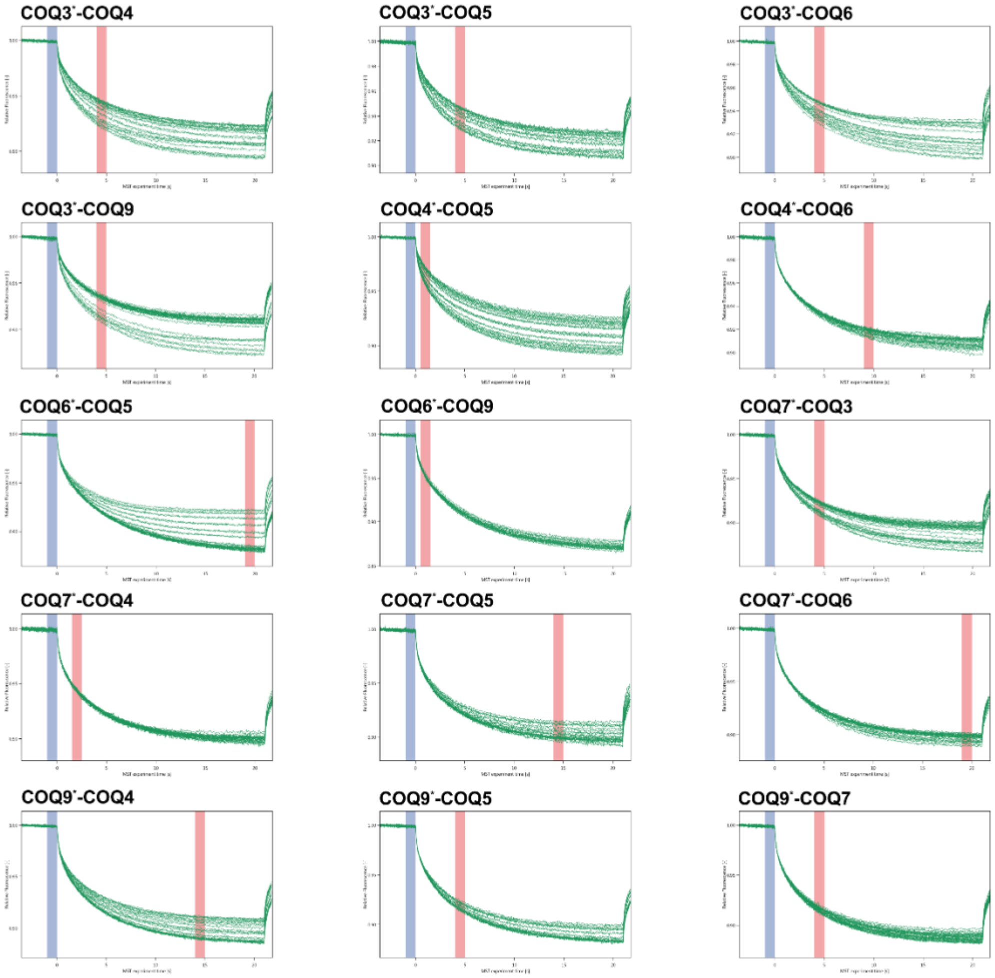
COQ3*-COQ4
COQ3*-COQ5
COQ3*-COQ6
COQ3*-COQ9
COQ4*-COQ5
COQ4*-COQ6
COQ6*-COQ5
COQ6*-COQ9
COQ7*-COQ3
COQ7*-COQ4
COQ7*-COQ5
COQ7*-COQ6
COQ9*-COQ4
COQ9*-COQ5
COQ9*-COQ7

**b**

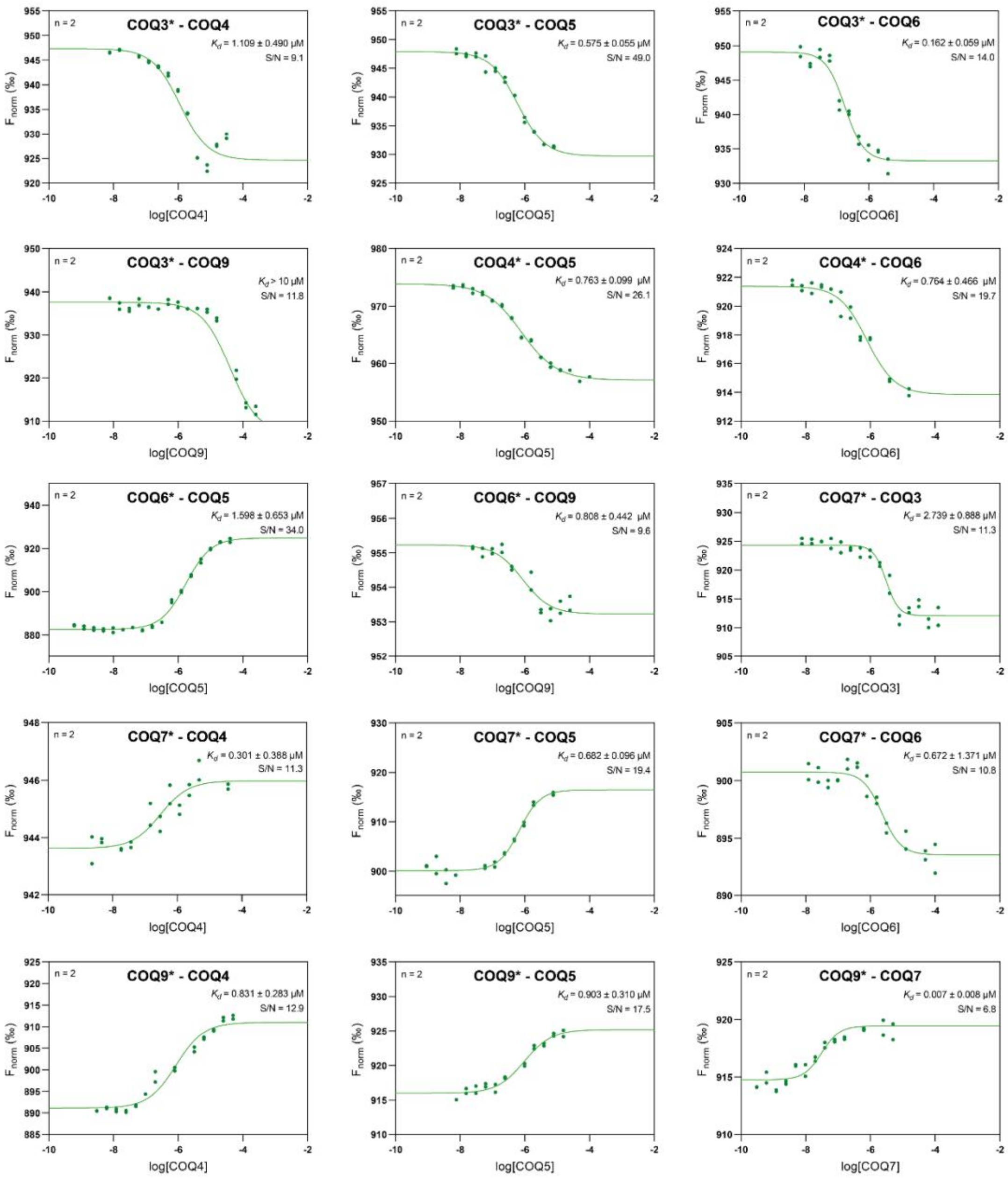


**Supplementary Figure 4. Protein-protein affinity measurements for COQ protein pairs using micro-scale thermophoresis and His-tag labelling strategy**. The His-tag fluorescently labelled COQ protein is marked with *. **a.** Experimental traces; the time used for plotting dose-response curves is highlighted in red, the reference signal used for normalization in blue. **b.** Dose-response curves. The labelled protein is marked with *; dissociation constants and signal to noise ratio (S/N) are reported. Data are reported as individual data point of n=2 independent experiments.

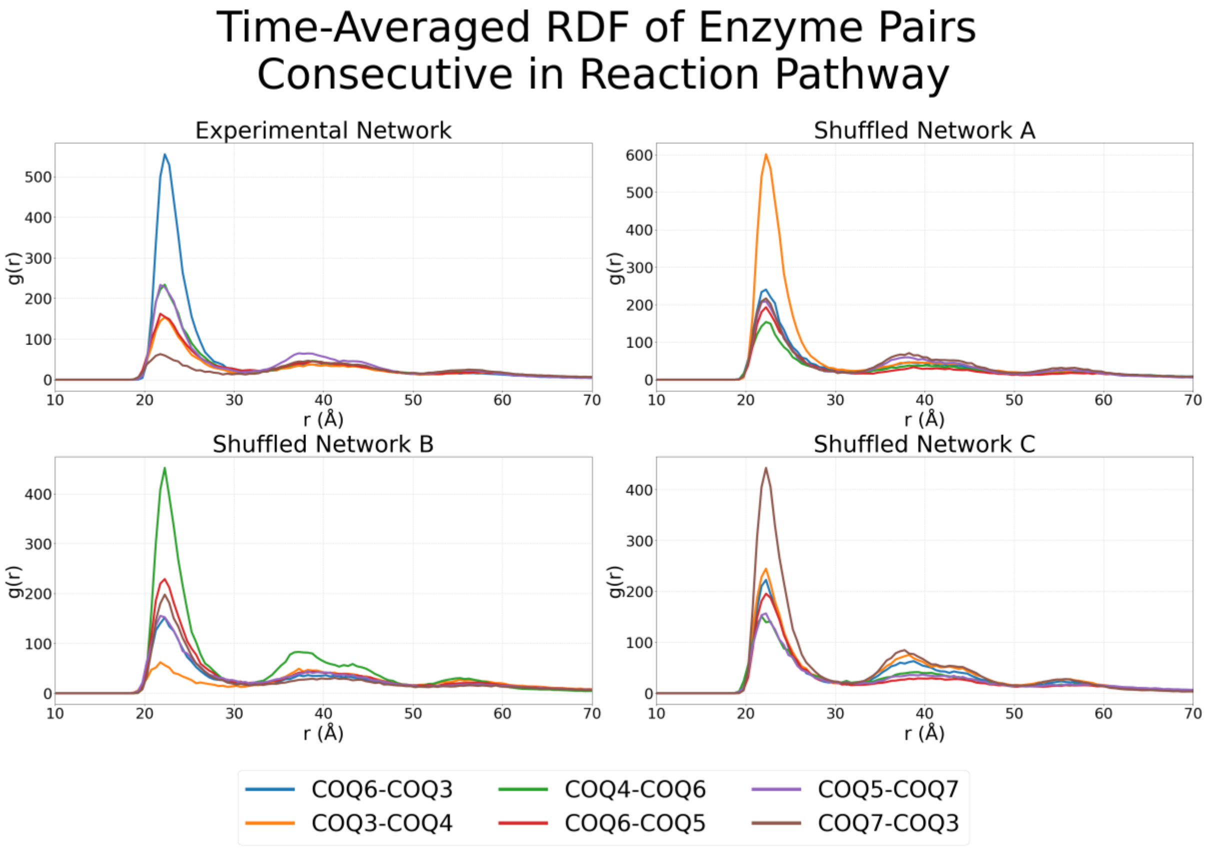


**Supplementary Figure 5. Interaction network determines metabolon fine structure.** Time-averaged radial distribution functions (RDFs) are shown for six enzyme pairs that catalyze consecutive steps in the pathway, comparing the unshuffled experimental network to node-shuffled network A, B, and C.

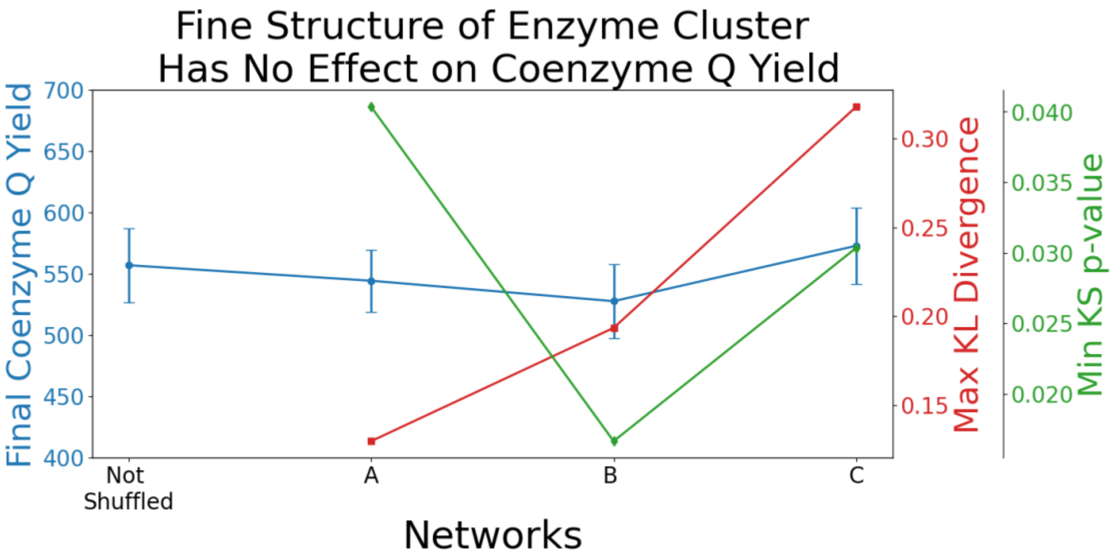


**Supplementary Figure 6. Fine structure of enzyme cluster has no effect on coenzyme Q yield.** Comparison of three node-shuffled networks (A, B, C) with the unshuffled network shows that, despite significant changes in RDF distributions, final coenzyme Q yield (blue) remains largely unaffected. The maximum KL divergence (red) reflects the greatest difference among the six enzyme pair RDFs between shuffled and unshuffled networks. The minimum KS p-value (green) indicates the most statistically significant difference in the six RDFs, with all three shuffled networks showing at least one RDF significantly different from the original ($p < 0.05$).

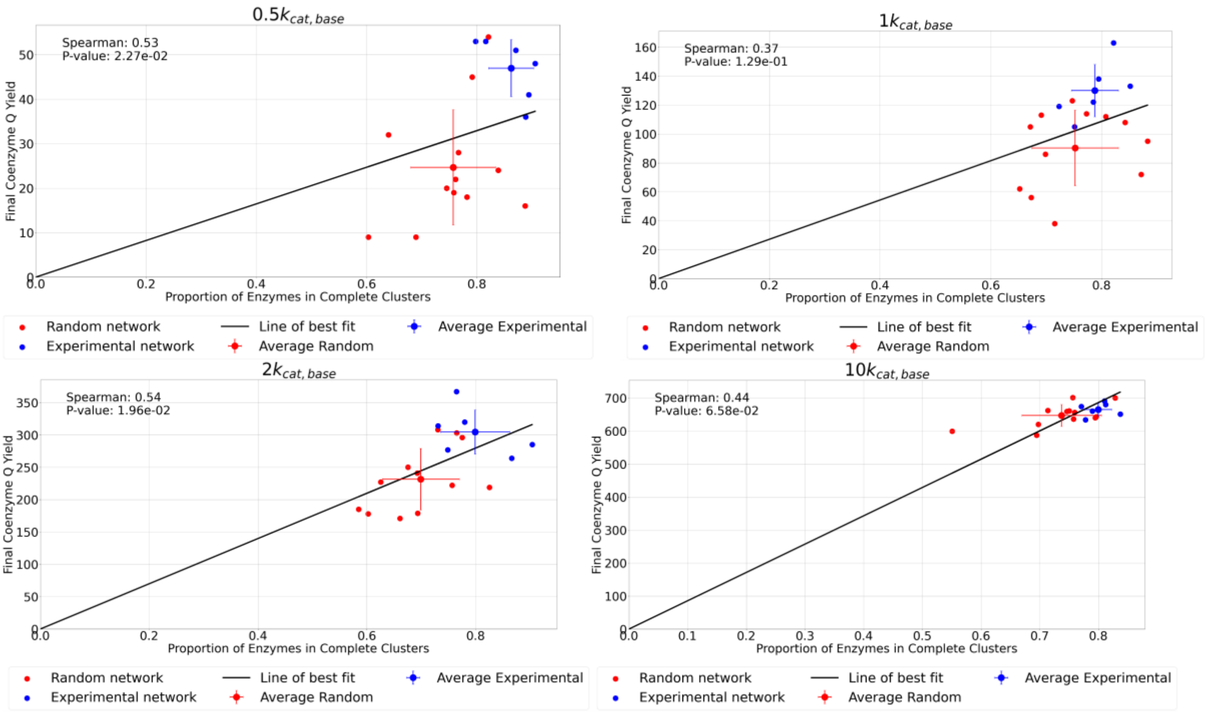


**Supplementary Figure 7. Correlation between coenzyme Q yield and the proportion of enzymes in complete clusters (containing at least one copy of COQ3-7) at increasing activity values.** Data points show individual simulations for experimental (blue) and randomized edge networks (red).

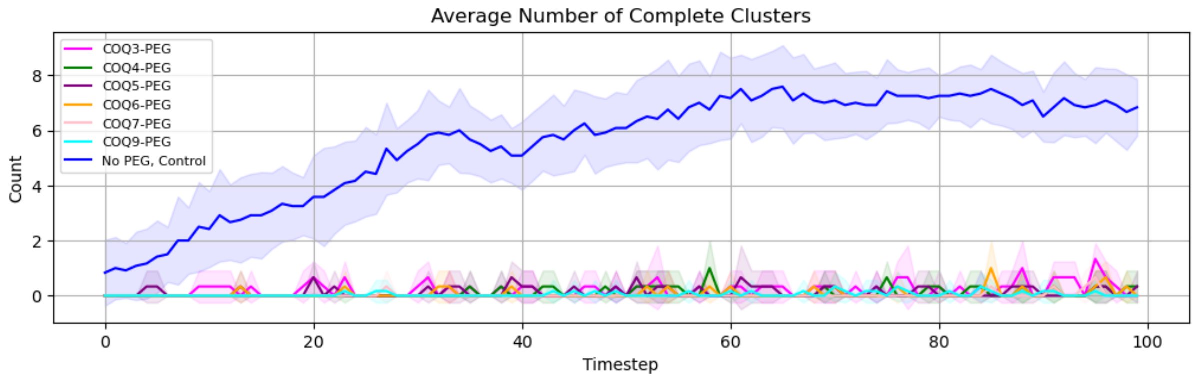


**Supplementary Figure 8. Progression of average number of complete clusters over time.** Error represents standard deviation (n=6 for control, n=3 for PEGylated systems).

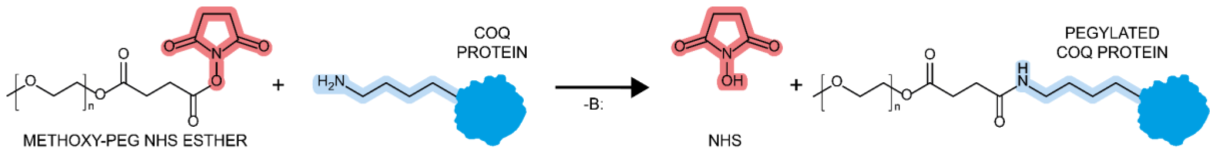


**Supplementary Figure 9. Reaction scheme of PEGylation of primary amino groups.**

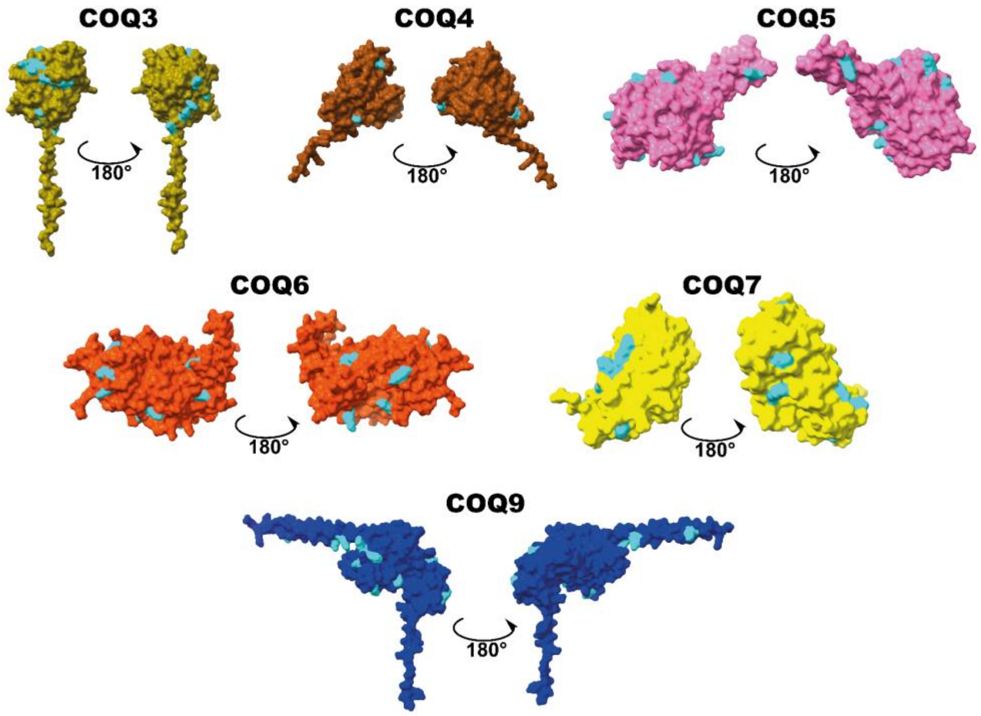


**Supplementary Figure 10. AlphaFold-3 predicted structures of COQs shown as surfaces**. Surface lysine residues are colored in cyan.

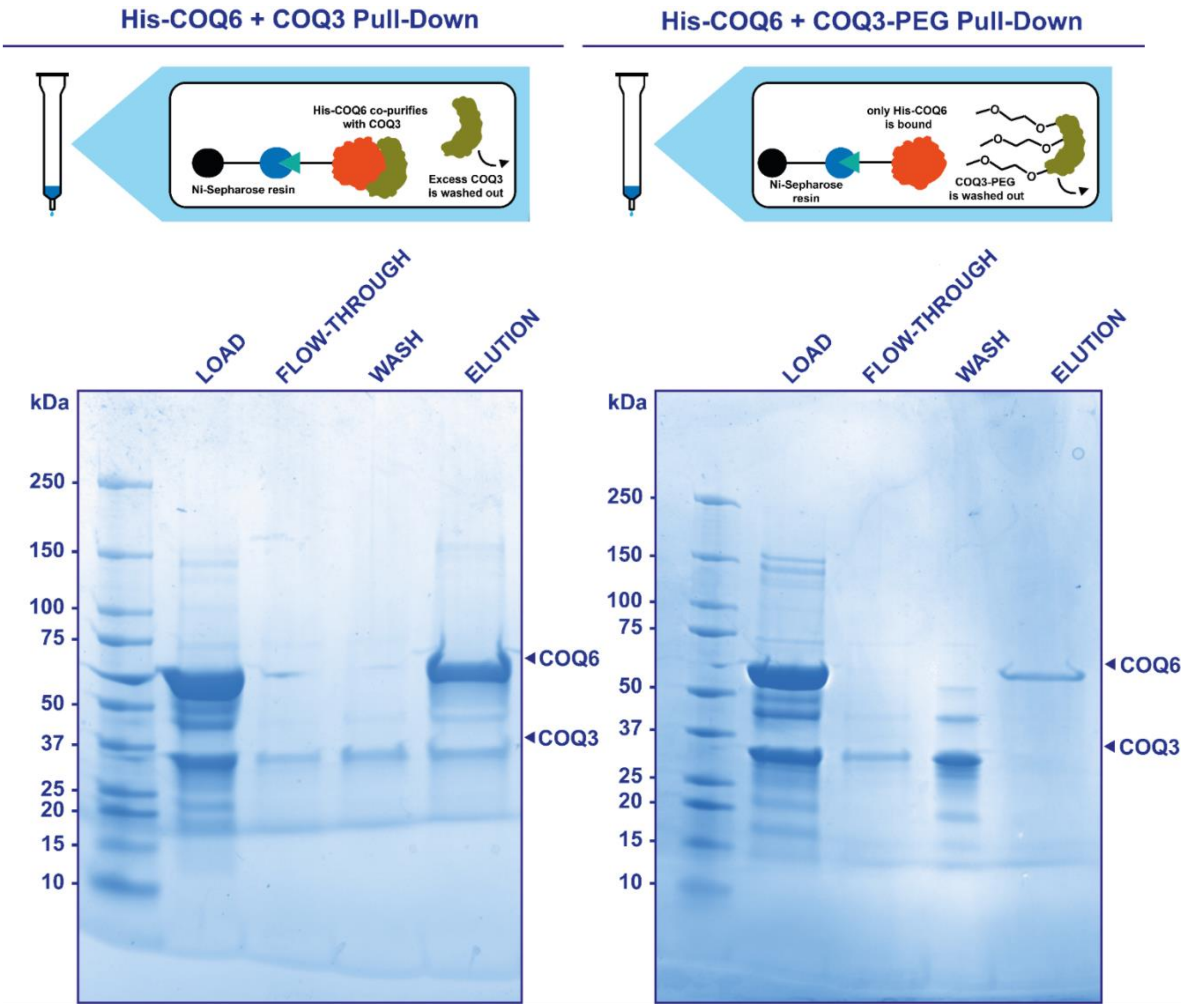


**Supplementary Figure 11. Pull-down assay demonstrates that PEGylation of COQ3 impairs its binding with COQ6.** The control assay with un-labelled COQ3 is shown on the left side, the assay with PEGylated COQ3 on the right. Reference molecular weight markers of COQ3 and COQ6 are outlined with an arrow on the right side of the gel. All replicates and uncropped gels are reported in Supplementary Fig. 20.

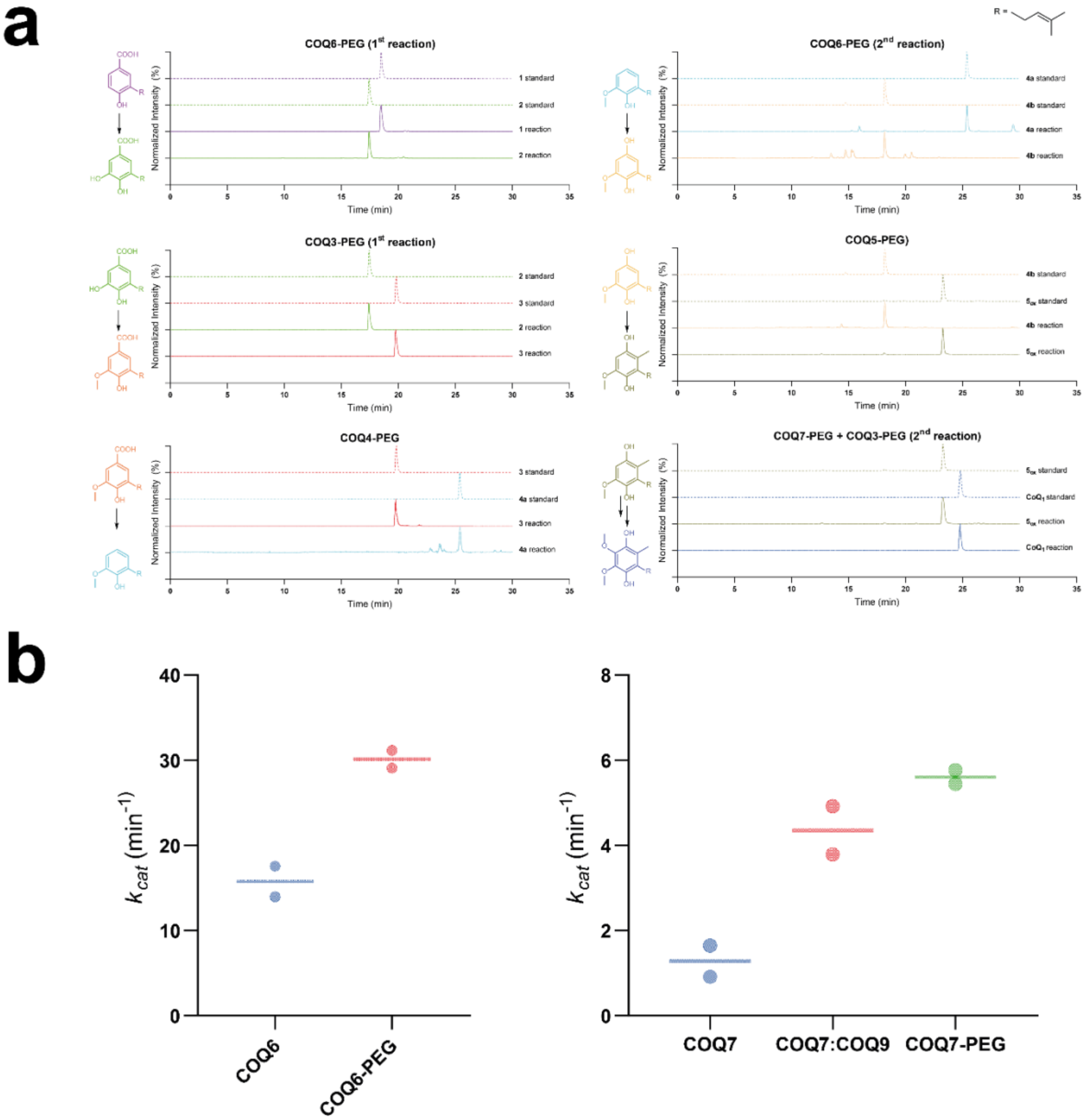


**Supplementary Figure 12. PEGylated COQs retain their enzymatic activities**. **a.** Normalized extracted ion chromatograms (XIC) of substrate and product for each COQ reaction. Chromatograms of standard injections are shown with dashed lines, while sample chromatograms with continuous lines. The reaction of COQ7 was assayed coupled to COQ3 due to lack of analytical standard for intermediate **$6_{ox}$**. **b.** NAD(P)H consumption activity of PEGylated COQ6 and COQ7. Individual data points from n=2 independent experiments are shown as dots, the mean value as dash.

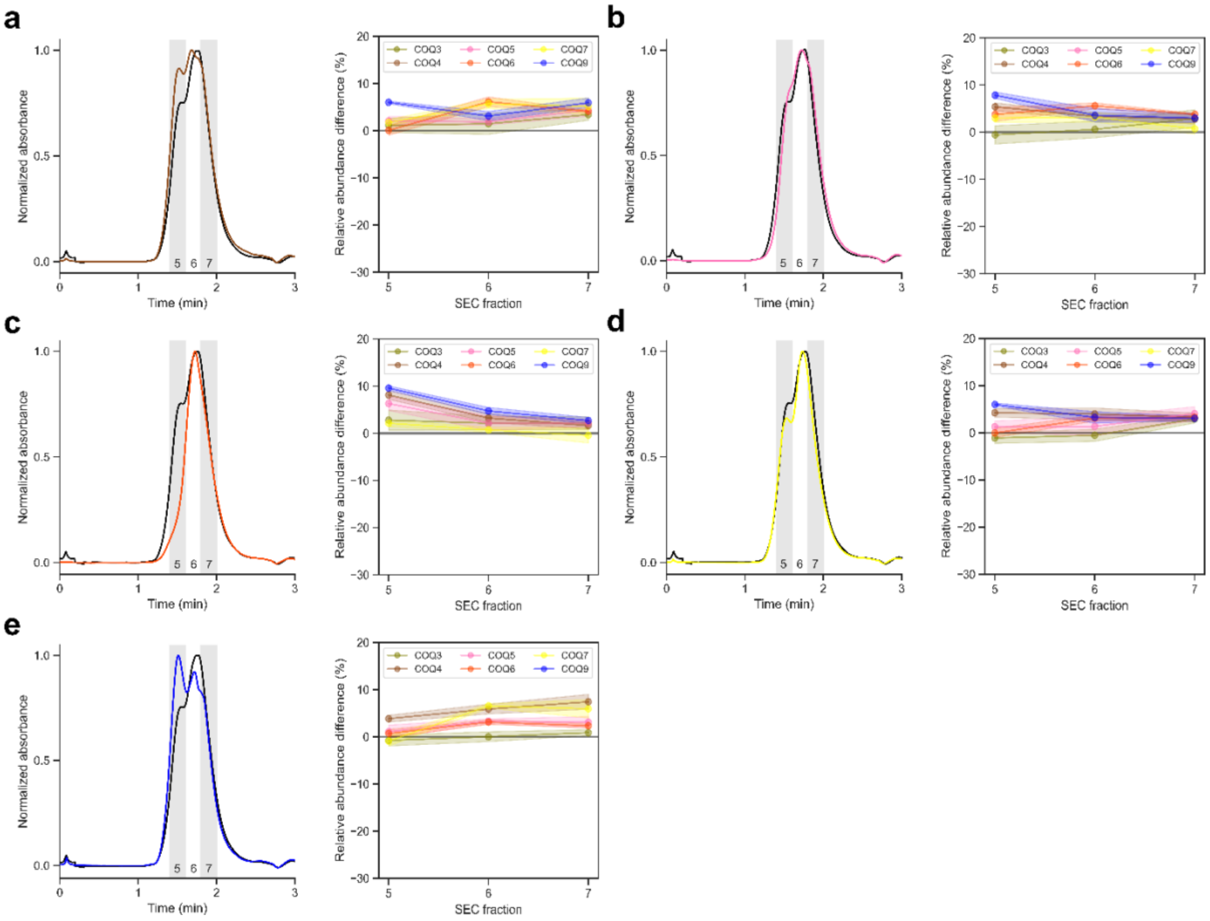


**Supplementary Figure 13. Analytical size-exclusion chromatography followed by bottom-up proteomics of the COQ metabolon depleted of individual proteins**. For each sample, the figure shows chromatograms of the intact COQ (black) and incomplete (colored) metabolons depleted of a certain COQ protein. The right panels highlight compositional differences between incomplete and intact COQ metabolons as quantified by bottom-up proteomics in n=3 independent experiments. Data are reported as mean and SD (shaded area). **a.** COQ metabolon without COQ4, **b.** COQ5, **c.** COQ6, **d.** COQ7, **e.** COQ9. SDS-PAGE analyses of each fraction is reported in Supplementary Fig. 19.

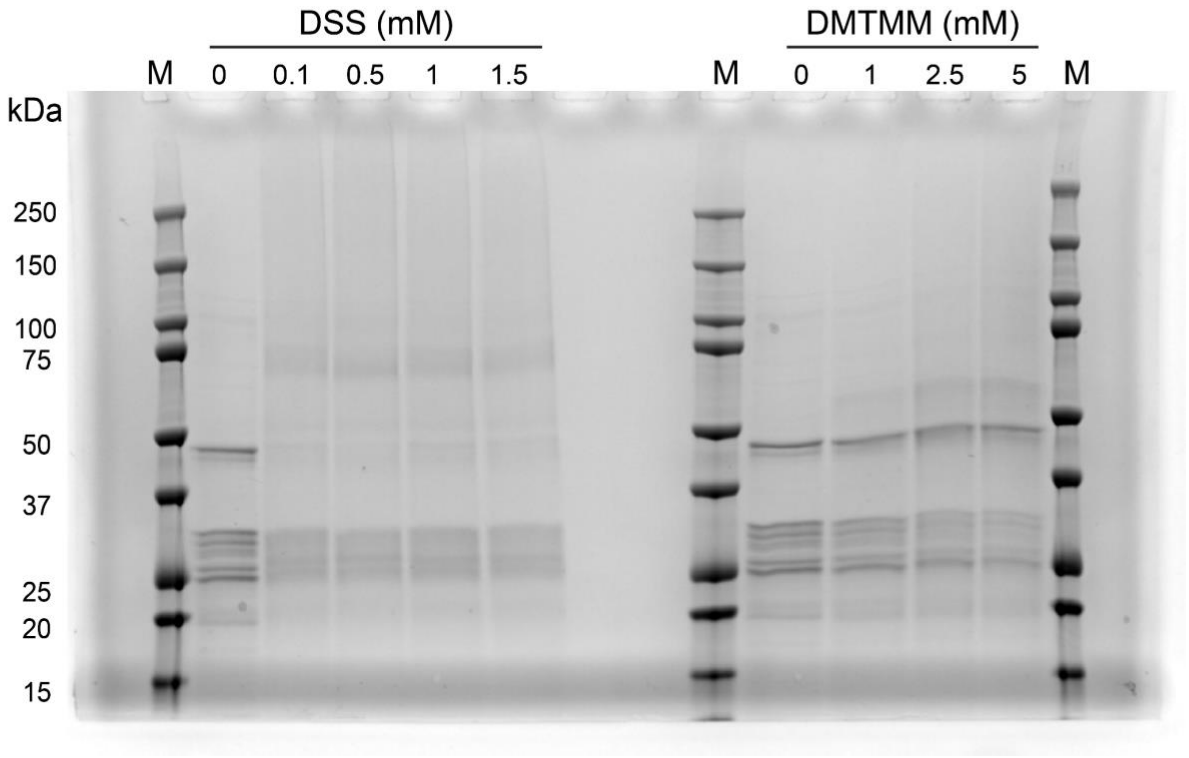


**Supplementary Figure 14. Uncropped reducing SDS-PAGE gel of cross-linking reaction optimization for COQ3-7, 9 sample** (Criterion XT Bis-Tris 4-12% Precast gel, Biorad). Each lane contains 2 µg of protein sample. DSS is disuccinimidyl suberate, DMTMM is 4-(4,6-dimethoxy-1,3,5-triazin-2-yl)-4-methyl-morpholinium chloride.

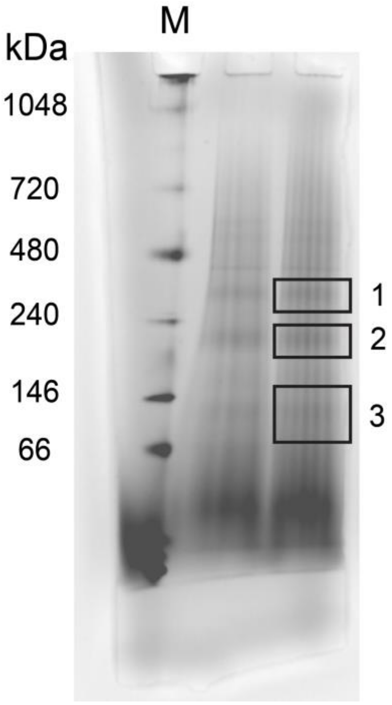


**Supplementary Figure 15. Blue native-PAGE of COQ3-7, 9 sample** (NativePAGE™ Bis-Tris Mini Protein Gel, 3 to 12%, 1.0 mm, Thermo Fisher)**.** The bands that were selected for in-gel XL-MS are highlighted and labeled (Supplementary Fig. 2). Each line contains 15 μg of protein sample.

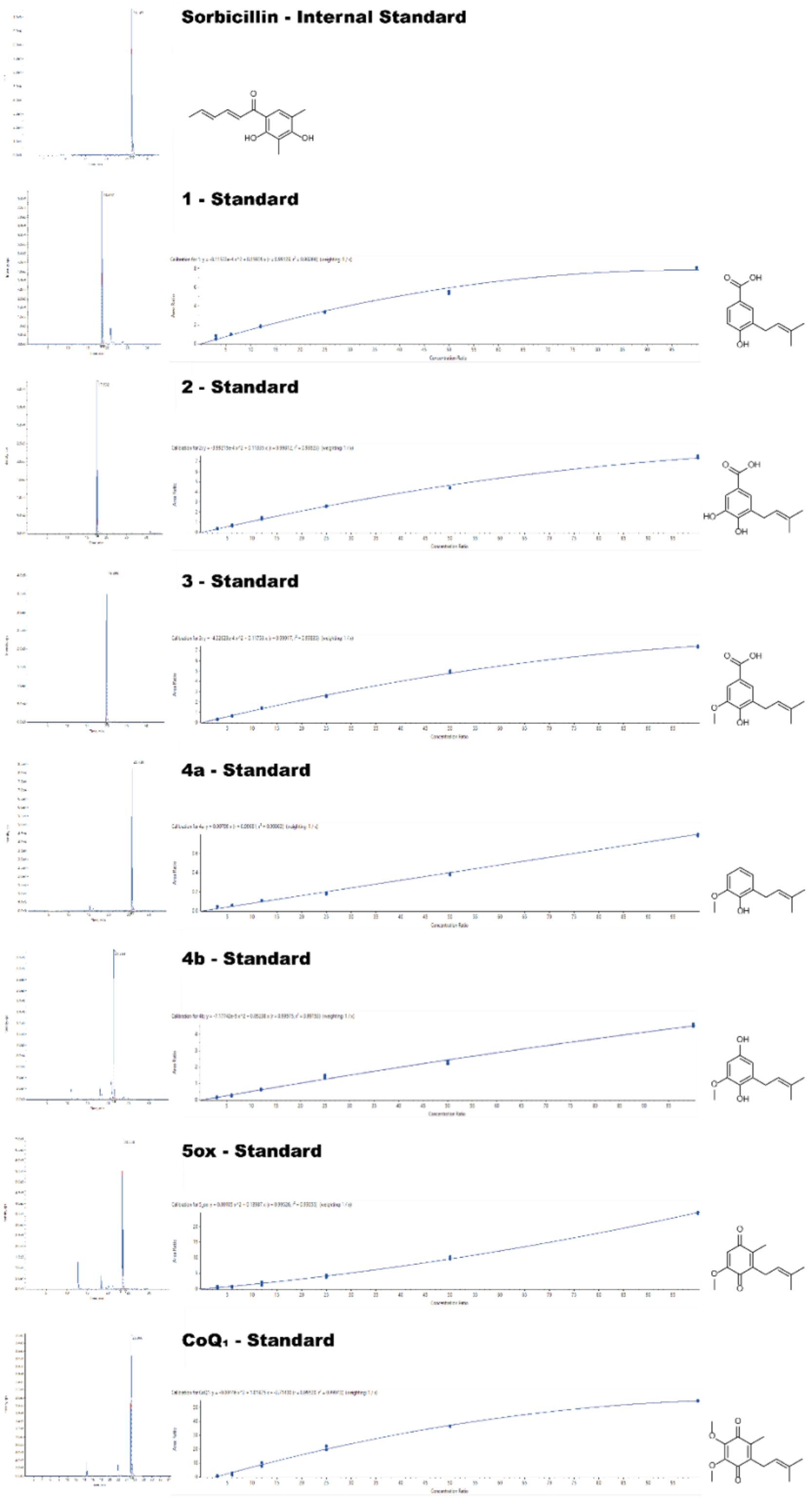


**Supplementary Figure 16**. **Automated peak integration and calibration curves of all analytes used for LC/MS quantitation experiments**. Peak integrations are shown on the left side for 50 µM analytical standards, except for the internal standard that was 1 µM. Quadratic nonlinear regression was applied to fit all calibrations, except for **4a** which was analyzed with linear regression.

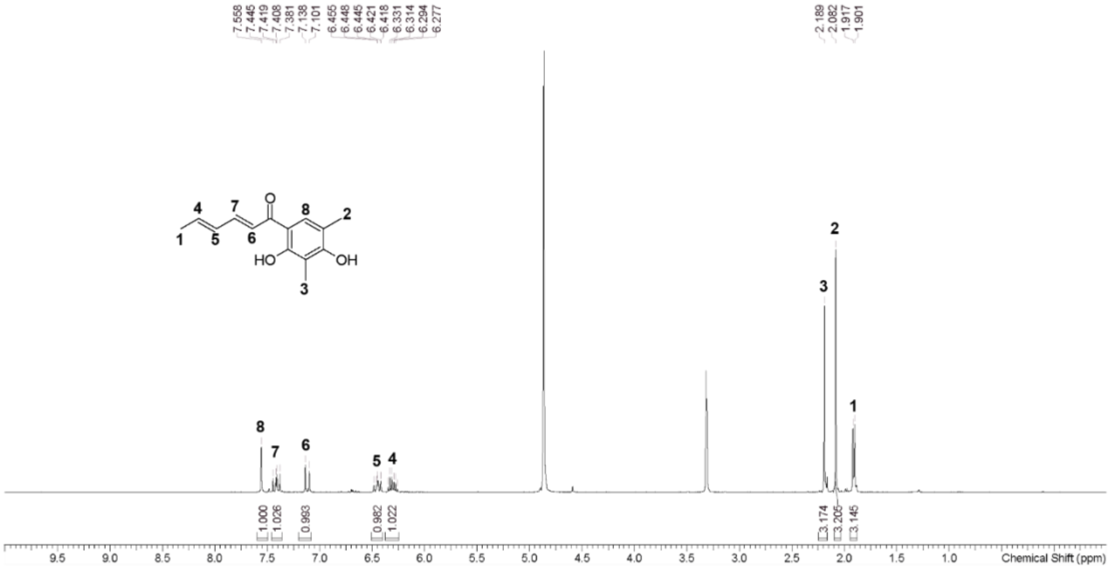

**Supplementary Figure 17. ¹H NMR spectra of the internal standard Sorbicillin.** Spectrum was acquired in MeOD. Assigned peaks are numbered at increased ppm values.

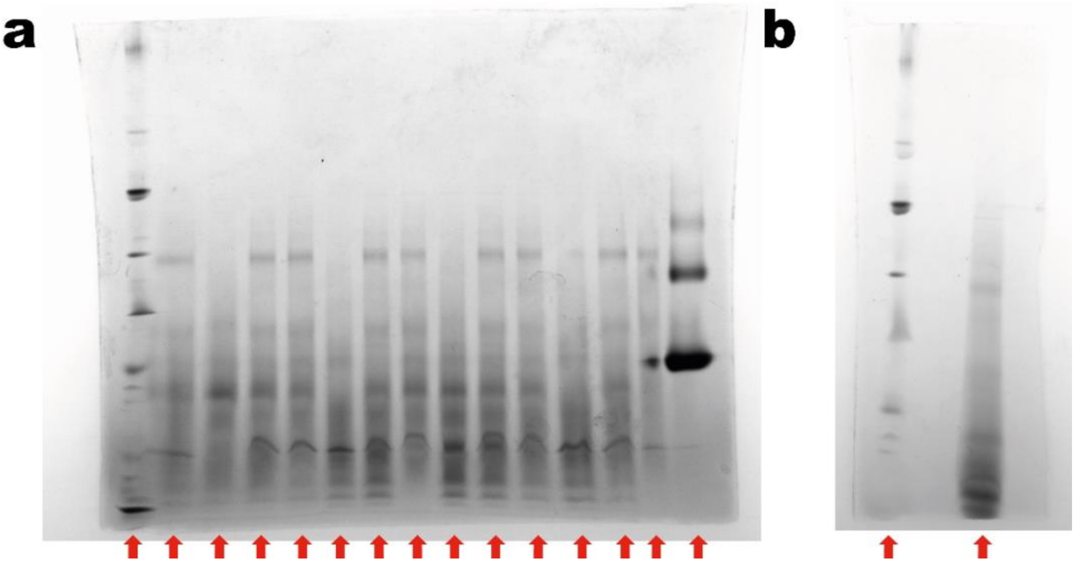

**Supplementary Figure 18**. **Uncropped Blue Native PAGE gels. Each lane reported in the main figures is marked by a red arrow**. **a.** Analysis of the COQ metabolon depleted of individual COQs or replaced with PEGylated COQs as shown in Fig. 6c. **b.** Analysis of the COQ metabolon with COQ4 replaced by an active site mutant as shown in Fig. 7b.

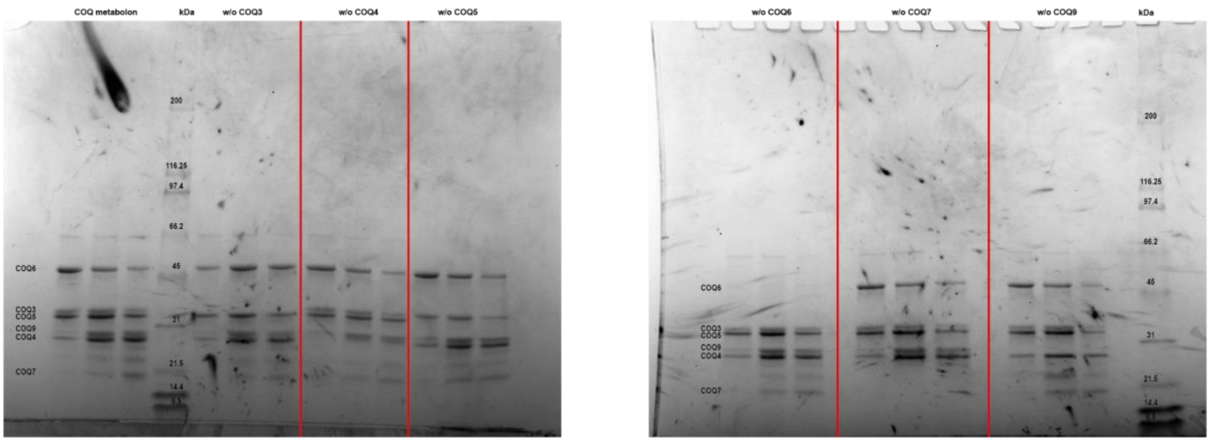


**Supplementary Figure 19. Uncropped SDS-PAGE analysis of fraction from chromatograms of Fig. 2d and Fig. 6d.** The gel is labelled with the reference chromatogram on top, with reference band height for each COQ protein on the left, and with MW on the ladder. For each chromatogram the three collected fractions were loaded in increasing order from left to right, each triplet is separated by a red line.

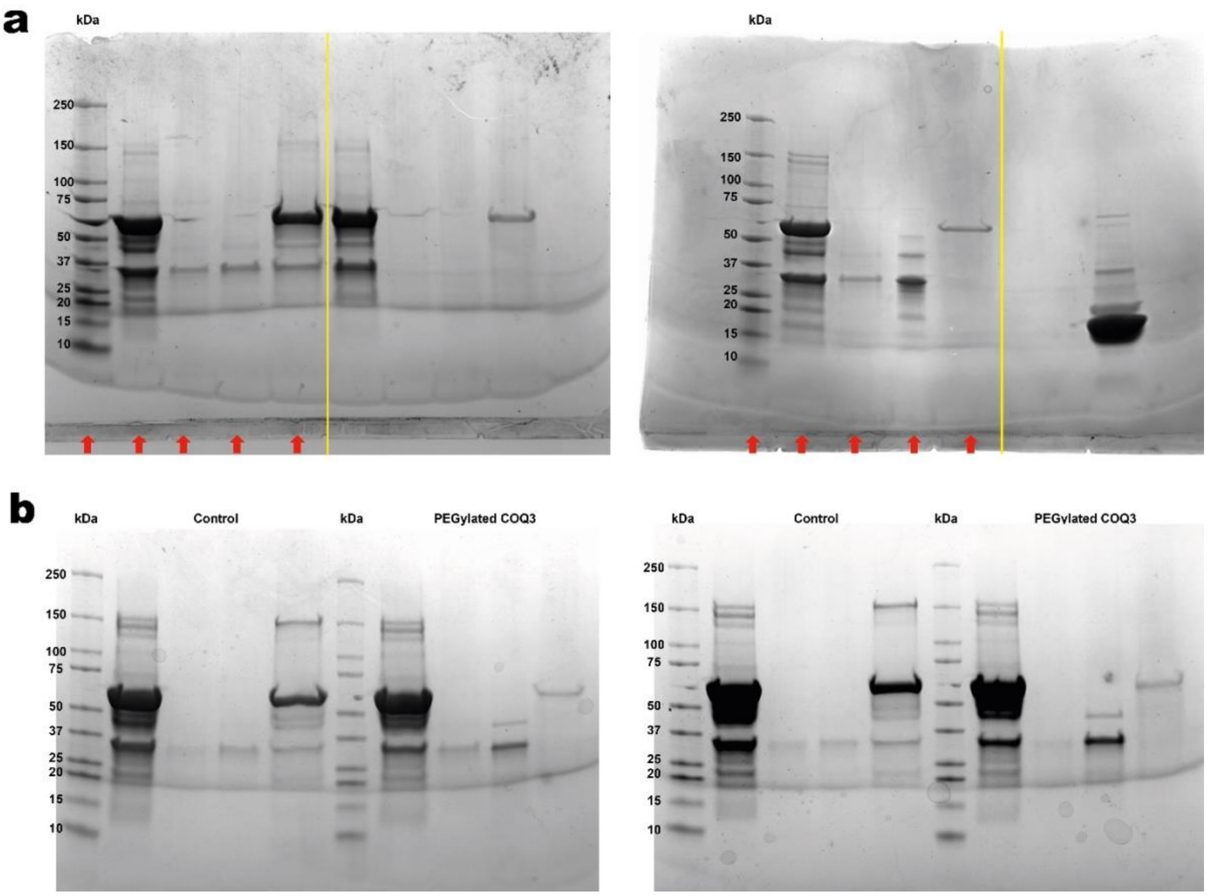


**Supplementary Figure 20. Uncropped SDS-PAGE analysis of the COQ6-COQ3 pull-down assay**. **a.** Uncropped gels from Supplementary Fig. 11. Each lane reported in the figure is marked by a red arrow, cropping is marked by yellow lines. The control experiment gel is shown on the left side panel, the one with COQ3 PEGylated on the right panel. **b.** Additional replicates of the pull-down assay. Each gel was loaded with ladder, load, flow-through, wash and elution, with the control experiment on the left side and the one with COQ3 PEGylated on the right side. Marker of the ladder is shown on the left, reference experiments on top.

### Supplementary Tables

| Band | Matched Protein | Coverage (%) | Matched Peptides |
|---|---|---|---|
| COQ Metabolon | COQ3 | 21.58 | 8 |
| | COQ4 | 26.23 | 6 |
| | COQ5 | 9.25 | 2 |
| | COQ6 | 21.58 | 8 |
| | COQ7 | 21.86 | 11 |
| | COQ9 | 22.59 | 8 |
| without COQ4 | COQ3 | 24.46 | 8 |
| | COQ6 | 14.58 | 8 |
| without COQ5 | COQ3 | 19.06 | 6 |
| | COQ6 | 17.31 | 10 |
| without COQ7 | COQ3 | 9.71 | 3 |
| | COQ6 | 16.4 | 8 |
| without COQ9 | COQ3 | 17.27 | 5 |
| | COQ4 | 62.3 | 38 |
| | COQ5 | 56.23 | 28 |
| | COQ6 | 50.11 | 60 |
| PEGylated COQ4 | COQ3 | 19.06 | 6 |
| | COQ6 | 17.08 | 9 |
| PEGylated COQ5 | COQ3 | 12.59 | 4 |
| | COQ6 | 13.67 | 7 |
| PEGylated COQ7 | COQ3 | 12.59 | 4 |
| | COQ6 | 19.82 | 12 |
| PEGylated COQ9 | COQ3 | 53.6 | 27 |
| | COQ4 | 60.66 | 33 |
| | COQ5 | 49.82 | 24 |
| | COQ6 | 47.38 | 55 |

**Supplementary Table 1. Peptide mapping analysis of the Blue Native PAGE gel from Fig. 6b.** The peptides were obtained from digestion of the band marked in the respective figures. The complete list of peptide sequences obtained from the analysis is provided as raw data.

| Band | Matched Protein | Coverage (%) | Matched Peptides |
|---|---|---|---|
| COQ Metabolon with COQ4mut | COQ3 | 22.66 | 8 |
| | COQ4 | 34.02 | 9 |
| | COQ5 | 11.03 | 6 |
| | COQ6 | 25.97 | 23 |
| | COQ7 | 20.22 | 6 |
| | COQ9 | 13.81 | 5 |

**Supplementary Table 2. Peptide mapping analysis of the Blue Native PAGE gel from Fig. 7b.** The peptides were obtained from digestion of the band marked in the respective figures. The complete list of peptide sequences obtained from the analysis is provided as raw data.
